# Gene-centric metagenomic analyses reveal microbiome functional insights into diseases

**DOI:** 10.1101/2025.09.29.679262

**Authors:** Shen Jin, Aurelie Cenier, Lea Eisenhard, Emilio Ríos, Daniela Wetzel, Woo Young Cho, Till R. Lesker, Tobias Eisen, Svenja Schorlemmer, Agnieszka Gaska, Tingting Zheng, Marilena Stamouli, Merianne Mohamad, Sudip Das, Vishal C. Patel, Till Strowig, Melanie Schirmer

## Abstract

The microbiome encodes millions of genes; however, understanding their role in human health remains challenging. Here, we developed MetaGEAR, a gene-centric analysis framework for metagenomic data. MetaGEAR constructs cohort-specific databases for efficient retrieval of gene annotations, abundances, and co-localization, while providing enhanced taxonomic resolution by integrating reference- and assembly-based approaches. This is combined with a Metagenomic Assembled Graph (MAGraph) capturing gene neighborhood information. Using MetaGEAR, we built a multi-cohort database comprising >33 million gene families to investigate microbiome functionality across 24 cohorts of inflammatory bowel disease, colorectal cancer, and healthy populations and identified disease signature genes. Furthermore, the MAGraph revealed mobile genetic elements acting as hubs for tetracycline resistance spreading among *Enterococcus*, *Streptococcus*, and *Veillonella* species. Also, a duplicated nitrate reduction operon in *Klebsiella pneumoniae* was linked to differential gene expression under stress and virulence-inducing conditions. In summary, gene-centric metagenomic analyses reveal important insights into microbiome functionality in diseases.

## Main

Millions of microbial genes can be profiled from metagenomic sequencing data from human cohort studies and subsequently summarized into non-redundant gene catalogs. Several gene catalogs are publicly available and based on large-scale metagenomic data analyses, including the Unified Human Gastrointestinal Protein (UHGP [1]) catalog and the Integrated Non-redundant Gene Catalog (IGC [2]). These serve as valuable reference databases for mapping-based functional profiling of new metagenomic data sets. However, two critical aspects are currently missing: (1) Gene neighborhood information is typically lost during the gene catalog construction and (2) current database infrastructures lack support for efficient and comprehensive extraction of abundance profiles, taxonomic classifications, and functional annotations.

Gene neighborhood information is critical for in-depth analyses of microbiome functionality. A distinctive feature of bacterial genomes is their operon structure, which are clusters of coregulated genes [3] [4]. Current graph-based tools, including Spacegraphcats [5] and STRONG [6], use gene neighborhood information based on assembly or co-assembly graphs. These tools are primarily designed to address strain heterogeneity and improve assembly completeness across multiple metagenomic samples [7] but lacks the scalability to produce comprehensive graph representations of the entire metagenomic cohort. While Spacegraphcats supports the extraction of neighboring genes, its scope is limited to small sample sets to generate co-assembly graphs but does not capture detailed information on genome organizations. Currently existing tools lack data structures that efficiently summarize and capture gene neighborhood information at scale, leaving essential features such as co-localization frequencies and cis/trans relationships largely inaccessible for microbiome functional analysis.

Scalability of gene catalogs has become a major bottleneck as cohorts are reaching hundreds to thousands of samples with increasing sequencing depth, resulting in tens of millions of gene families. Querying a gene of interest from gene abundance profiles with hundreds of billions of records requires an efficient database infrastructure. The large majority of gene families are present in only a few samples, resulting in many zero entries in the abundance tables. This data sparsity poses challenges for traditional SQL-based systems, which typically model abundance profiles as dense tables with gene families as rows and samples as columns. In contrast, NoSQL databases offer flexible data models better suited to sparse and heterogeneous data, making them promising data structures for large gene catalogs.

To address these challenges, we developed a gene-centric analysis framework, termed **MetaG**enomic **E**xploration **A**nd **R**esearch (MetaGEAR), that includes three major advancements: Firstly, we built an end-to-end pipeline that integrates reference- and assembly-based methods to generate detailed functional profiles, while increasing species-level annotations by ∼10% on average. These profiles are stored in a NoSQL database, enabling rapid queries within seconds across billions of records. Secondly, we applied this pipeline to 9,053 stool metagenomes from inflammatory bowel disease (IBD), colorectal cancer (CRC) and healthy individuals, generating a human gut gene catalog with over 33 million microbial gene families with consistent taxonomic and functional annotations. This comprehensive data resource enabled the identification of disease signature genes, including an IBD-enriched *Hungatella hathewayi* trehalase gene, which breaks down anti-inflammatory trehalose and thereby potentially contributes to a pro-inflammatory gut environment. Lastly, by generating a Metagenomic Assembled Graph (termed MAGraph) we facilitate the exploration of gene neighborhoods. Using the MAGraph, we identified novel insights into host-associations of mobile genetic element (MGE) carrying antibiotic resistance genes and strain-specific operon structures associated with disease. Overall, the gene-centric framework of MetaGEAR enables scalable, functional exploration of the microbiome facilitating new insights into its role in human health.

## Results

### Overview of the MetaGEAR analysis framework

The first component of our gene-centric framework, MetaGEAR, is a standardised metagenomic analysis pipeline to generate functional microbiome profiles for any given metagenomic dataset. Its main output is a NoSQL database with a non-redundant gene catalog, which includes information on gene abundances, functional annotations, and taxonomic assignments. NoSQL databases, such as MongoDB, offer flexible data models with greater scalability for massive data storage [8] [9], making them promising frameworks for managing metagenomic data resources. Specifically, MongoDB leverages sparse data by storing only non-null fields, thereby reducing storage requirements while improving query efficiency. We used MongoDB to enable fast and efficient gene queries that can typically be executed within seconds (**Figure 1a**). The developed Application Programming Interfaces (APIs) allow the user to query for abundance information, functional and taxonomic annotations, gene sequences, and genomic locations (including contig and strand information) across cohort samples, using gene/protein sequences and functional domains as input (**Figure 1b**). Additionally, assembly-based approaches, including Metagenomic Species Pangenomes (MSPs) generated by MSPminer [10], often suffer from low detection sensitivity and limited taxonomic resolution. For MetaGEAR, we developed an integrated reference- and assembly-based approach, where we match MSPs to MetaPhlAn species through abundance correlations (**Figure 1c**), leveraging their complementary strengths and generating detailed functional profiles with improved taxonomic annotations. The second part of MetaGEAR is a unified multi-cohort database, which combines information from 24 publicly available metagenomic studies and facilitates the investigation of microbiome functionality in health and disease across cohorts (**Figure 1d**). For this, we applied our MetaGEAR pipeline and re-analyzed 9,053 metagenomic stool samples from IBD, CRC, and healthy individuals (**Table 1**). Gene families from each cohort were aggregated to construct a multi-cohort gene catalog (MCGC) comprising over 33 million non-redundant microbial gene families. The NoSQL data model enables efficient queries and access to consistent taxonomic annotations and abundance profiles across all cohorts. We also constructed a multi-cohort protein catalog (MCPC) for efficient queries of protein sequences and functional domains and a metadata database (MCMD) that includes coherent age, gender, and disease information for all cohorts. The last component of our framework is the MAGraph, a novel data structure that uses the information from the NoSQL database to capture and visualize gene neighborhood information from the metagenomic assemblies (**Figure 1e**). In the MAGraph, nodes represent gene families and edges indicate their co-localization on assembled contigs, allowing the reconstruction of gene neighborhoods across diverse microbial genomes from the gut microbial communities. Users can search the MAGraph for genes of interest by sequence or functional domains and extract the respective gene neighborhood graph. This neighborhood graph includes information on inter-gene distances, strand orientation (cis/trans), and co-occurrence frequencies across cohort samples (**Figure 1e**). The MAGraph is a powerful data structure for visualising and uncovering microbiome functionality, such as host identification of mobile genetic elements and strain-specific bacterial operon structures or genomic context.

**Figure 1:**
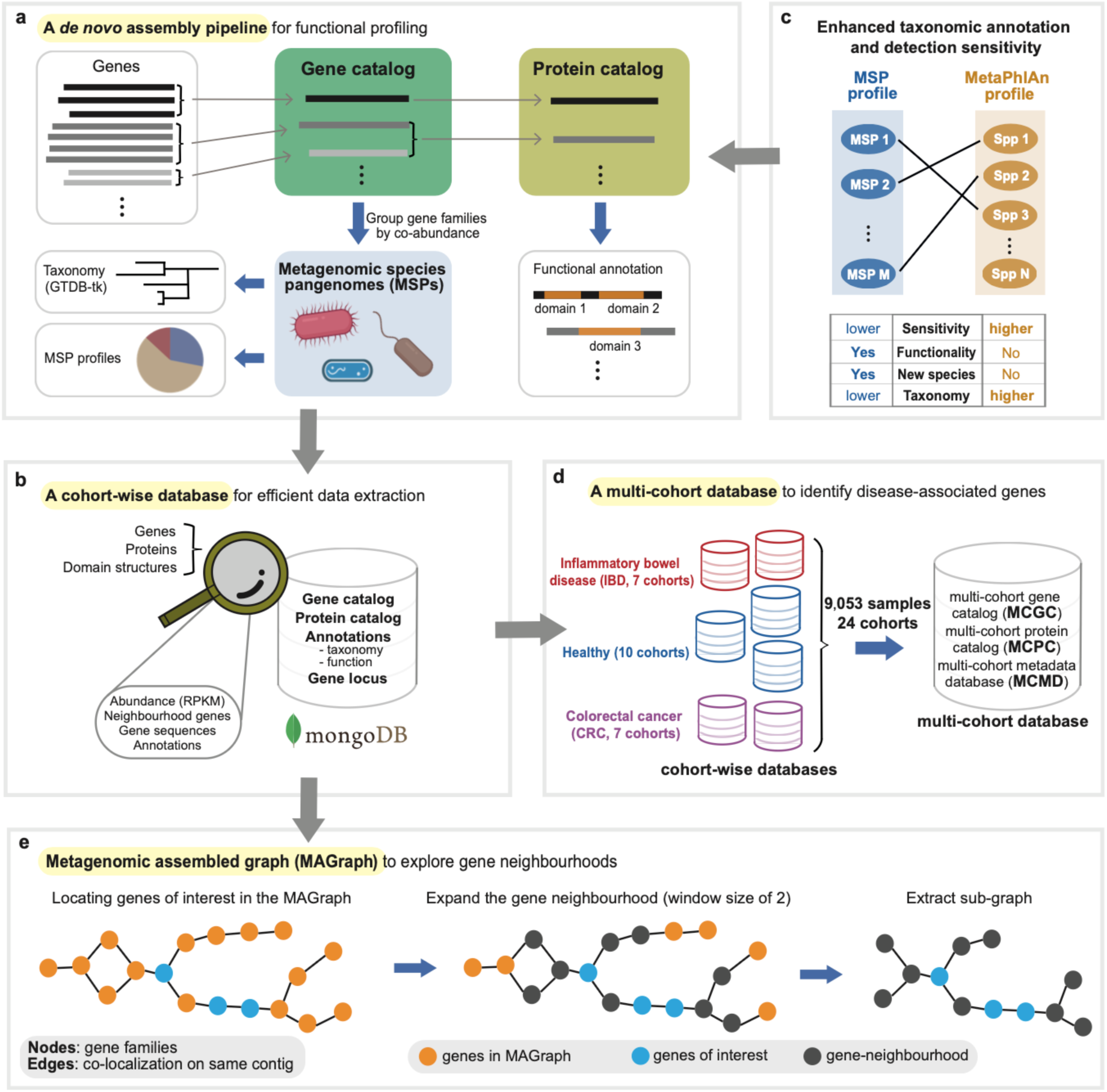
MetaGEAR overview: a gene-centric approach for metagenomic analyses. (**a)** First, a *de novo* assembly pipeline is used to generate functional profiles. A non-redundant gene catalog is generated for a given metagenomic dataset, where microbial genes are grouped at species-level based on sequence similarity (identity >95%, coverage >90%). Species-level gene families are then further grouped into metagenomic species pangenomes (MSPs) based on co-abundance patterns. Taxonomic annotations of MSPs are inferred using GTDB-tk and MSP abundance profiles are calculated as median abundance of their core genes. The gene catalog is then further grouped into a protein catalog based on amino acid similarity (identity >90%, coverage >80%) and Pfam domains are annotated for representative protein sequences. **(b)** A cohort-wise NoSQL database (MongoDB) is built to facilitate efficient data extraction for the functional profiles, which includes gene and protein sequences, abundances, taxonomic and functional annotations, and gene localization information. **(c)** Integration of reference-based (MetaPhlAn3) and assembly-based (MSPs) analyses results in species-level microbiome profiles with (1) increased detection sensitivity and taxonomic resolution, (2) gene profiles for each species, and (3) profiles of new species without a known reference genome. **(d)** A multi-cohort database facilitates the identification of disease-associated genes across cohorts. For this, we processed 24 cohorts (9,053 metagenomic stool samples) with MetaGEAR, focusing on inflammatory bowel disease (IBD), colorectal cancer (CRC), and healthy individuals. The resulting cohort-wise databases were combined into a multi-cohort database, where cohort-specific gene families were grouped into a multi-cohort gene catalog (MCGC) and subsequently summarised into a multi-cohort protein catalog (MCPC). The metadata from individual studies was curated and stored in a multi-cohort metadata database (MCMD). **(e)** Construction of a metagenomic assembled graph (MAGraph) to capture and analyze gene neighborhoods. Each node represents a gene family and edges connect adjacent gene families located on the same contig. The graph structure facilitates searching for gene(s) of interest (i.e. blue nodes) and extraction of neighboring genes (i.e. gray nodes, here: window size = 2) as a sub-graph.

### Integrating reference- and assembly-based profiling approaches to enhance taxonomic annotations of metagenomic species pangenomes

This integrative analysis approach is based on the comparison of a species’ abundance based on reference-based profiling (MetaPhlAn3 [11]) and its corresponding assembled metagenomic species pangenome (MSP [10]), which should be linearly correlated. We confirmed this using species annotated by both approaches. For example, the abundance of *Bacteroides dorei* inferred by MetaPhlAn showed a highly significant linear correlation with the MSP annotated as *B. dorei* by GTDB-tk [12] [13], which was consistent across all cohorts (**Figure S1a**, r^2^>0.95 and p-value<10^-180^ for all cohorts). Importantly, MetaPhlAn showed high sensitivity and was able to reliably detect low-abundant occurrences of *B. dorei* that were missing in the assembly-based profiles (highlighted samples along the x-axis, **Figure S1a**). Our integrative approach takes advantage of these abundance correlations, resulting in several improvements: Firstly, MSP taxonomic annotations were substantially increased, where we identified annotations for an additional 10% of the MSP-profiled communities with this approach (**Figure S1b**). Likewise, *de-novo* assembled gene profiles can be linked to species identified by reference-based taxonomic profiles, thus providing insights into their functional potential. On average corresponding MSPs were identified for 80% of the MetaPhlAn-profiled communities (**Figure S1c**). Interestingly, the abundance of unannotated MSPs explained the majority of unannotated reads in MetaPhlAn profiles (**Figure S1d**) and unannotated MSPs accounted on average for ∼50% of the microbial community per sample (**Figure S1c**). Notably, the accuracy of the MSP taxonomic assignment was consistently high (**Figure S1e**, mean accuracy = 97.3%±5.1%). Here, accuracy was evaluated based on comparing high-confidence MSPs and their inferred MetaPhlAn match (see method section for details). Overall, these analyses indicate that integrating reference- and assembly-based approaches for taxonomic profiling of metagenomic sequencing data combines the complementary advantages of these approaches resulting in improved taxonomic resolution while revealing information about gene content of the detected species.

Interestingly, this hybrid approach can also be used to estimate the detection limit of MSPs, which can provide a guideline for choosing an optimal sequencing depth. Based on a subset of confidently assigned MSP-MetaPhlAn species pairs, we inferred an empirical estimation for the detection limit for each cohort (details in the method section). The inferred detection limit was highly consistent across all cohorts, where species were rarely detected if fewer than 10,000 reads mapped to the corresponding MSP (empirical detection likelihood <25%, details in the method section). Conversely, MSPs of species with more than 100,000 reads were confidently detected (empirical detection likelihood >75%, **Figure S2a**). For the sequencing depth, this implies that 10 million reads (after quality control) are required per sample in order to reliably assemble species with a relative abundance of at least 1%.

In current metagenomic studies many opportunistic pathogens are commonly missed by assembly-based approaches, such as *Veillonella*, *Streptococcus,* and *Klebsiella spp.*, which can be present at low abundance in early or mild stages of disease (**Figure S2b**). Our hybrid-based approach has the potential to greatly improve the detection sensitivity for these species, resulting in assembly-based profiles of lowly-abundant species. For example, using an assembly-based approach alone, the IBD-associated species *Veillonella parvula* [14] was only detected in 27.6% of samples from patients in remission and showed no significant changes in mild ulcerative colitis [UC] (IBD_Elinav_2022 cohort, p-value=0.19). However, using our hybrid approach we were able to detect *V. parvula* in 78.6% of remission samples (**Figure S2c**), revealing a significant increase in patients with mild UC compared to controls (Wilcoxon, p-value=0.02, **Figure S2d**). In summary, our integrative analysis approach improves taxonomic resolution, in particular for lowly-abundant species, which can reveal important microbiome signals in early disease stages.

### Identifying IBD and CRC signature genes with a comprehensive multi-cohort database

A major challenge in microbiome studies is to identify disease-enriched genes that are consistently implicated across different cohorts. To address this challenge, we performed a meta-analysis and applied our MetaGEAR pipeline to 24 publicly available metagenomic cohorts (n_total_=9,053 stool samples) with a focus on IBD (7 cohorts), CRC (7 cohorts) and healthy populations (10 cohorts) (**Figure 2a**). All cohorts were coherently processed with our pipeline and then combined into a multi-cohort database, consisting of three major components: a multi-cohort gene catalog (MCGC), a multi-cohort protein catalog (MCPC), and a multi-cohort metadata database (MCMD). The MCGC contains >33 million non-redundant species-level gene families (grouped at 95% similarity and 90% coverage [15] [2] [16] [17]). Of these gene families, seven million were commonly detected and present in at least 2 cohorts (**Figure 2b**). By merging the taxonomic information based on the individual cohorts, we were able to significantly improve taxonomic annotations for the multi-cohort database, resulting in 3.4 million MCGC gene families with species-level annotations. These originate from 28.4 million cohort-wise gene families, among which 52% (14.9 million) were originally unannotated. Currently, this multi-cohort database represents the largest human gut microbiome gene catalog (**Figure S2e**). Importantly, our database structure facilitates the efficient retrieval of gene abundance profiles, taxonomic and functional annotations, gene sequences, and gene locations. Our database includes gene-abundance information for more than 300 billion entries (33 million gene families across >9 thousand samples) in addition to information for 707 million *de novo* assembled gene sequences.

**Figure 2:**
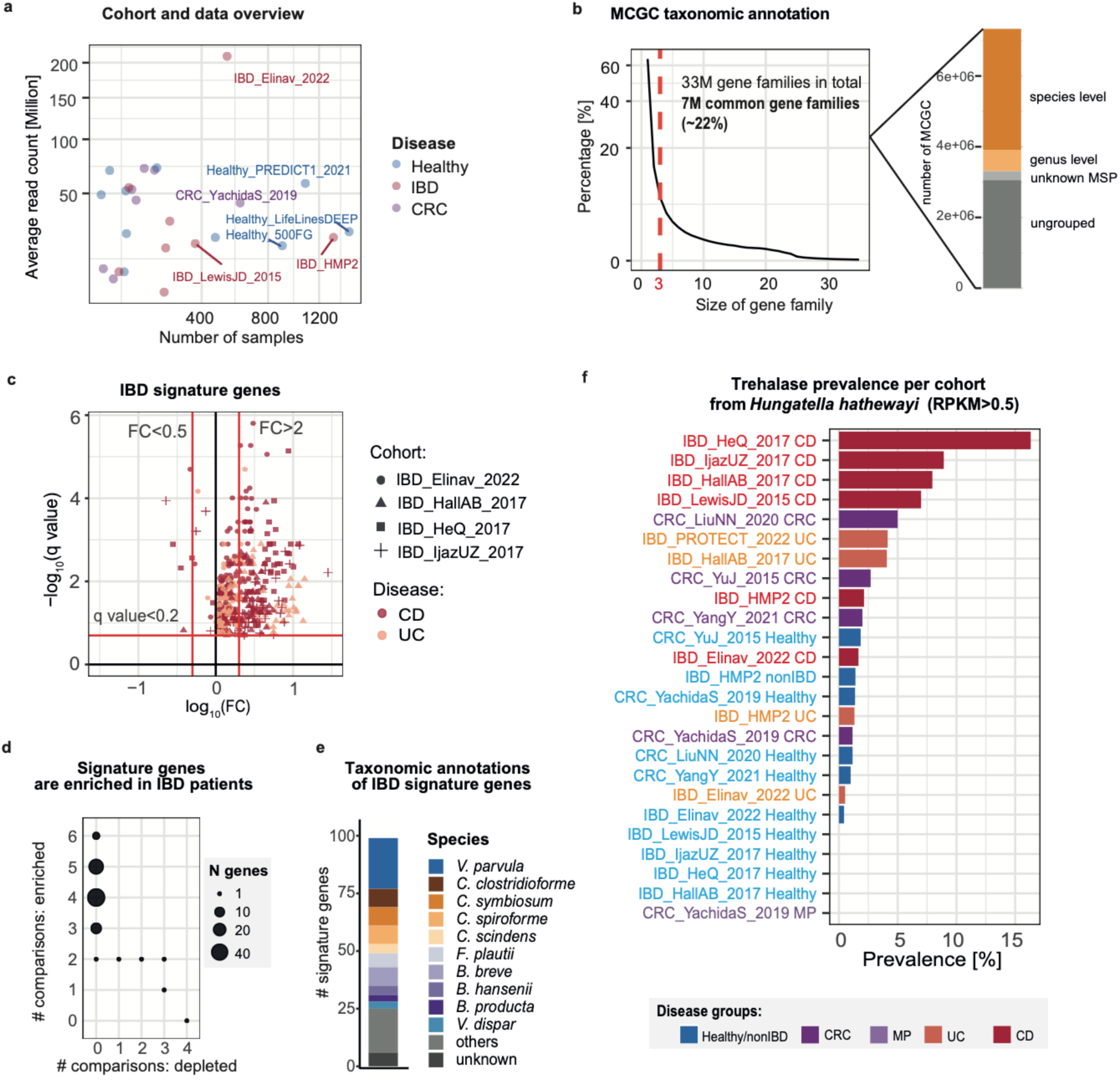
Identification of Inflammatory bowel disease (IBD) signature genes based on the multi-cohort database. **(a)** Data overview for cohorts including total number of samples (x-axis) for each cohort and average read count per sample (y-axis). Colors indicate disease focus. **(b)** Overview of the taxonomic annotations of the Multi-Cohort Gene Catalog (MCGC), a non-redundant gene catalog constructed by grouping representative sequences from cohort-level gene families. **Left**: The distribution of the sizes of MCGC gene families. We identified approximately seven million (7 M) common MCGC gene families, defined as those that contain at least three cohort-level gene families (size ≥ 3). **Right**: Taxonomic annotations for common MCGC gene families, where the y-axis indicates the number of gene families in the MCGC and color shows the taxonomic annotation level. **(c)** IBD gene enrichment including q-values (Wilcoxon, −log10 transformed, BH correction) versus mean fold change (FC, log10 transformed, disease vs control). Color indicates the disease group and red lines highlight thresholds. **(d)** IBD signature genes are consistently enriched in IBD patients across different studies. The plot shows the number of comparisons supporting a given gene as IBD enriched (y-axis) vs. IBD depleted (x-axis). The point size indicates the number of gene families in each group. **(e)** Taxonomic annotations of the IBD signature genes with the number of MCGC clusters (y-axis). The color indicates the corresponding taxonomic annotations at species-level. **(f)** Prevalence comparison of trehalase from *Hungatella hathewayi* across different cohorts (RPKM>0.5 per cohort disease group) showing an enrichment of this gene in IBD. Color indicates disease group (colorectal cancer -CRC, mixed polyp - MP, ulcerative colitis - UC, Crohn’s disease - CD).

We used this multi-cohort database to identify IBD and CRC signature genes. These are genes that are consistently enriched in patients across all cohorts and characteristic for the disease, i.e. these genes were predominantly or exclusively detected in the disease of interest. To identify IBD signature genes, we first selected genes that were present in at least three IBD cohorts (without distinguishing between IBD patients and controls) but mostly absent in other cohorts (e.g. healthy populations or CRC cohorts). These pre-selected genes were subsequently analyzed within each IBD cohort to determine if their abundance was significantly increased in IBD patients compared to controls. Most of the genes that were characteristic for the IBD cohorts were indeed enriched in IBD patients (**Figure 2c**). Analogously, we inferred CRC signature genes (**Figure S2f**). Notably, these disease signature genes represent robust gene signals, consistently enriched across multiple cohorts (**Figure 2d+S2g**). IBD signature genes were predominantly contributed by *Veillonella*, *Clostridium,* and *Blautia spp.* (**Figure 2e**), many of which have been previously implicated in IBD [14] [18] [19] [20]. Importantly, our analysis implicates specific genes for these IBD-associated species. For example, trehalase activity was the most enriched Gene Ontology (GO) term among the IBD signature genes and predominantly contributed by *Hungatella hathewayi* (**Table 1**). In particular in Crohn’s disease patients trehalase prevalence was consistently increased across multiple studies, while the gene was rarely detected in healthy individuals (**Figure 2f**). Trehalase is a microbial enzyme that converts trehalose, a common dietary compound [21] with anti-inflammatory effects [22], into glucose. This suggests that *H. hathewayi* may contribute to a pro-inflammatory disease state by reducing trehalose levels in IBD patients.

### Using gene neighborhood information from MAGraphs to reveal gene hubs in the human gut microbiome

Genomic organization plays a key role for the function and regulation of prokaryotic genes, but this information is currently missing in existing large-scale gene catalogs of human gut microbiomes [2] [1]. Here, we developed a graph-based structure, termed MAGraph, which encodes gene neighborhood information inferred from metagenomic assemblies. In the MAGraph, each node represents a gene family from the metagenomic cohort and adjacent genes that are located on the same contig are connected by an edge (**Figure 1e**). The MAGraph facilitates the analysis and visualisation of gene neighborhoods, enabling functional insights including disease-associated patterns.

New biological insights can already be inferred from the general properties of the MAGraphs (for all 24 cohorts) based on the node-degree distribution (**Figure 3a**). Most nodes were of degree two (46%), indicating that linear structures dominate the MAGraph. However, about 1% of the nodes exhibited high centrality (node degree >10). These hub nodes were enriched for DNA integration activities, such as transposase activity, DNA strand exchange activity and DNA binding (**Figure 3a**, right panel). Furthermore, the majority of the nodes were organized as one connected component comprising 61% ± 12% of the total nodes in the graphs (**Figure S2h**). This suggests that while MAGraphs are predominantly composed of linear structures, DNA integrases serve as hubs connecting diverse microbial species within the graph. While previous studies have suggested that horizontal gene transfer events are common in the gut microbiome [23] [24] [25] [26], our results emphasize that the gut microbiome functions as a cohesive entity where sharing of genetic material through mobile elements strongly shapes each individual microbial community.

**Figure 3:**
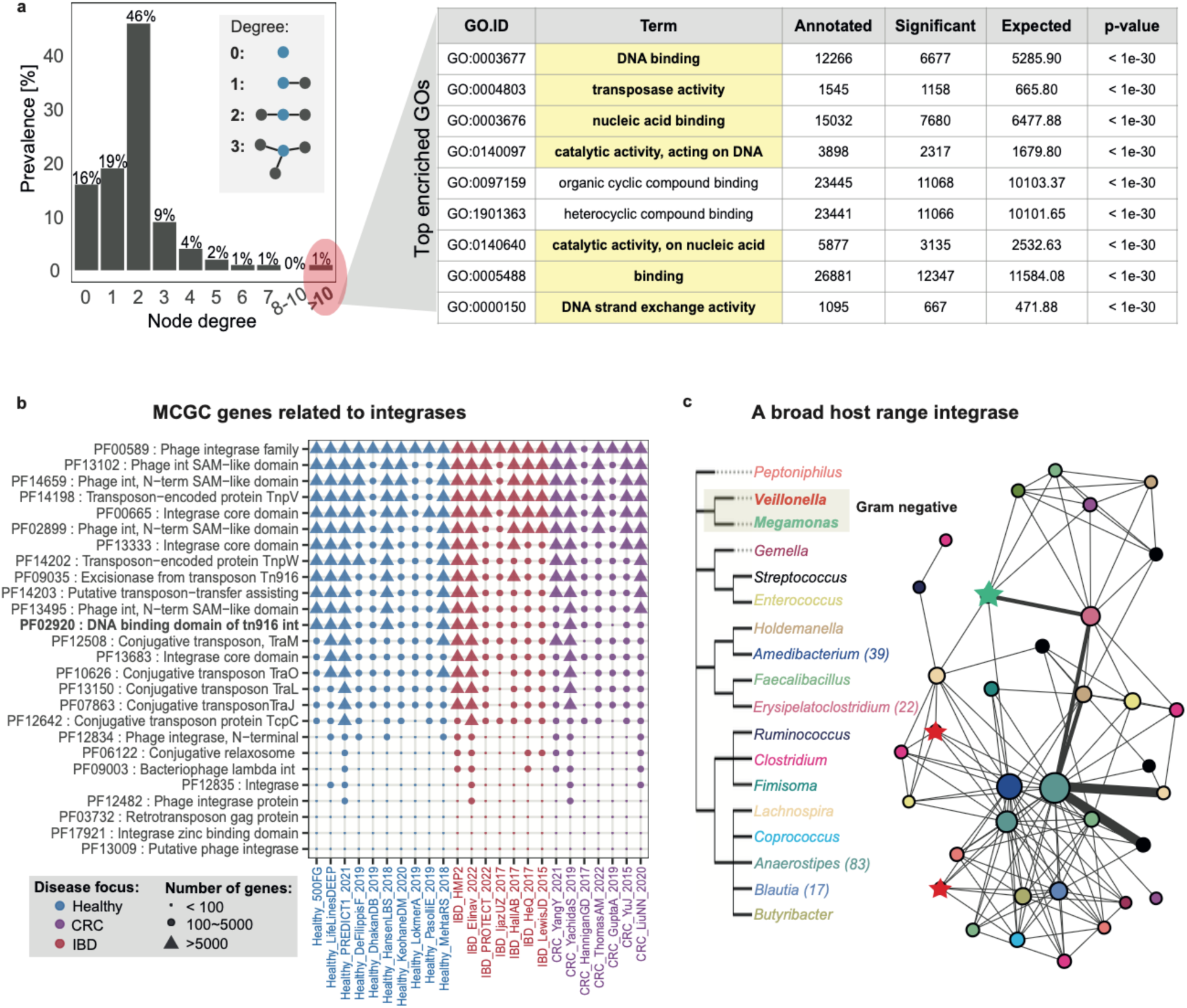
Mobile genetic elements act as hubs connecting diverse species in the MAGraph. **(a)** MAGraph statistics: **Left**: Node degree distribution (calculated per cohort and merged), indicating node degree (x-axis) and prevalence across all cohorts (y-axis). **Right**: GO enrichment analysis for highly connected nodes (degree>10, background: 100,000 randomly selected MCGC genes). The most enriched GOs terms are related to horizontal gene transfer (highlighted in yellow), such as DNA binding, transposase activity, and DNA exchange activity. **(b)** Profiling integrase-related genes in the multi-cohort gene catalog (MCGC). Each row indicates an integrase-related Pfam domain and each column contains information about the number of genes (shape) per cohort containing this domain. Color indicates disease focus. **(c)** Focusing on one specific MCGC gene family, a Tn196-like integrase, that was shared by multiple species from diverse genera. **Left**: Phylogenetic tree listing all genera containing this integrase, including two Gram-negative and 16 Gram-positive genera. The four most common genera are *Anaerostipes*, *Amedibacterium*, *Erysipelatoclostridium,* and *Blautia* (numbers indicate frequencies). **Right**: A network of all species sharing this integrase gene, where each node represents a species (colored by genus). The size of a node indicates how frequently the integrase gene was observed as part of the species’ genome. Both Gram-positive (round nodes) and Gram-negative bacteria (star nodes) were found to carry this integrase gene. Edges connect species in which the integrase appears on the same MAGraph (within the same cohort, n = 24). The thickness of an edge reflects how often the integrase is observed in the MAGraph with an edge linking these two species (the thicker the edge, the higher the frequency).

### Identification of a mobile element carrying tetracycline resistance with a broad host range

We further investigated the hub nodes involving integrase genes and uncovered a wide diversity of integrase families within all 24 metagenomic cohorts. Grouping integrase gene families by their PFAM domains revealed that phage-associated integrase hub nodes contained the highest number of gene families, followed by transposon-related integrases (**Figure 3b**). Next, we inferred the bacterial host range for each integrase gene family by examining the taxonomic annotations of their neighbouring genes in the MAGraphs across all cohorts. The most diverse host range and highest prevalence was observed for a highly conserved Tn916-like integrase gene family (average sequence identity >99%), which was assembled independently within all 24 metagenomic cohorts (1,221 out of 9,053 samples) and shared between 18 different genera within the phylum *Bacillota* (**Figure 3c**). While this gene family was most commonly detected in commensal Gram-positive bacteria, including *Anaerostipes* and *Amedibacterium spp.,* it was also found in opportunistic pathogenic *Negativicutes* species (Gram-negative), including *Veillonella* and *Megamonas*.

The high prevalence of this Tn916-like integrase enabled us to investigate mutation rates at each position using multiple sequence alignments of the identified 1,221 gene sequences (**Figure 4a**). Several sites exhibited high mutation rates leading to non-synonymous changes. Notably, one mutation in the 30th amino acid led to a change from hydrophobic isoleucine to hydrophilic threonine. This polymorphism occurred within the integrase DNA-binding domain potentially altering the integration site of the mobile element and facilitating its ability to infect diverse bacterial hosts. Importantly, this showcase highlights the potential of leveraging the growing number of genes assembled across metagenomic samples to identify key sites, which are evolutionarily conserved and potentially functionally important.

**Figure 4:**
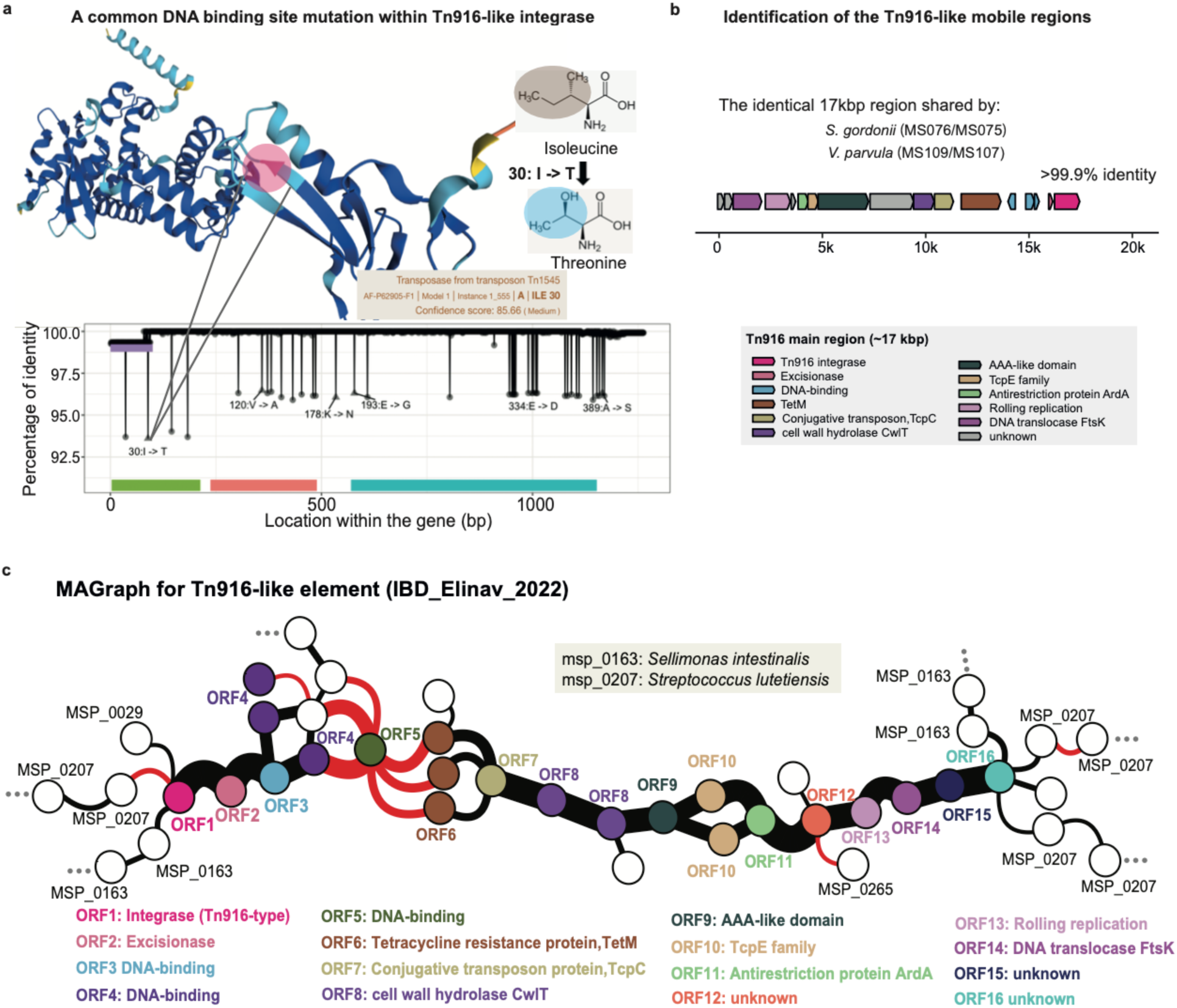
Identification of a 17 kbp broad-host-range mobile element harboring tetracycline resistance. **(a)** A common mutation occurs within the DNA binding domain for this Tn916-like integrase. **Upper part**: Predicted integrase structure (alphaFold2) where the mutated site within the DNA binding domain is highlighted (30: I -> T) turning isoleucine (hydrophobic) to threonine (hydrophilic). **Lower part**: Per-site mutation rates are plotted with mutation positions on the x-axis and percentage identity on the y-axis. All missense mutations are annotated. A color bar below the plot indicates domain annotations: green and red represent two DNA-binding domains, while blue refers to the core integrase domain. **(b)** This 17kbp-long Tn916-like integrase was also identified in clinical isolates of *Streptococcus gordonii* and *Veillonella parvula* (sequence identity>99.9%). Functional annotations for all genes are indicated by color. **(c)** A representative MAGraph for the IBD_Elinav_2022 cohort showing the subgraph of the Tn916-like mobile genetic element (MGE). Two MSPs, *Sellimonas intestinalis* (msp_0163) and *Streptococcus lutetiensis* (msp_0207) shared this MGE. Node colors represent matches to gene queries from the MGE, while white indicates no match to any query. Edge color reflects the strand orientation of adjacent gene families (black = same strand, red = opposite strand). Edge thickness corresponds to the frequency of colocalization between adjacent gene families, while edge curvature indicates the average distance between adjacent genes (straight lines: mean distance <50 bp, curved lines: mean distance >50bp).

To further investigate the functionality of this mobile element, we screened our clinical bacterial strain collection for isolates containing this integrase. Interestingly, this integrase gene was part of an identical 17 kbp region shared between *Streptococcus gordonii* and *Veillonella parvula* isolates (**Figure 4b**, >99.9% identity). Detailed annotation of this region revealed a Tn916-like element carrying the tetracycline resistance gene *tetM*, which encodes a ribosome protection protein (RPP) that confers tetracycline resistance by binding to the ribosome and displacing the drug from its binding site [27]. We further confirmed this in the IBD_Elinav_2022 cohort [28], where this mobile element was present, commonly shared by different species and a carrier of *tetM* (i.e. *Sellimonas intestinalis* and *Streptococcus lutetiensis*, **Figure 4c**, ORF6 annotated as *tetM*). The widespread presence of this *tetM*-carrying mobile genetic element among gut microbes suggests an important role in tetracycline resistance spreading.

We next confirmed that tetracycline resistance is associated with the presence of this MGE using clinical isolates with and without the Tn916-like element, including *Streptococcus parasanguinis*, *Streptococcus gordonii*, *Veillonella parvula,* and *Enterococcus faecium* isolates (average nucleotide identity >99.99%). Ten isolates from these four species were included in a tetracycline resistance assay, comparing the Minimal Inhibitory Concentration (MIC) of the strains carrying the mobile element against the strains lacking it (**Figure 5a**). The MIC was determined using E-test TC for all strains after 24 hours of tetracycline incubation confirming that strains without the Tn916-TetM mobile element were tetracycline-sensitive (*V. parvula* MS055, MS168 and MS209, *S. parasanguinis* MS080, and *S. gordonii* MS214, MIC < 1 ug/mL), while strains carrying the mobile element exhibited resistance (*V. parvula* MS107, 32ug/mL; *S. parasanguinis* MS081, 24ug/mL and MS082, 8ug/mL; *S. gordonii* MS075, 24ug/mL; *E. faecium* MS154, 256ug/mL, **Figure 5b+c**). This confirms that this MGE is present in a broad range of bacterial hosts and suggests that it is potentially a major factor in tetracycline resistance spreading among clinical isolates.

**Figure 5:**
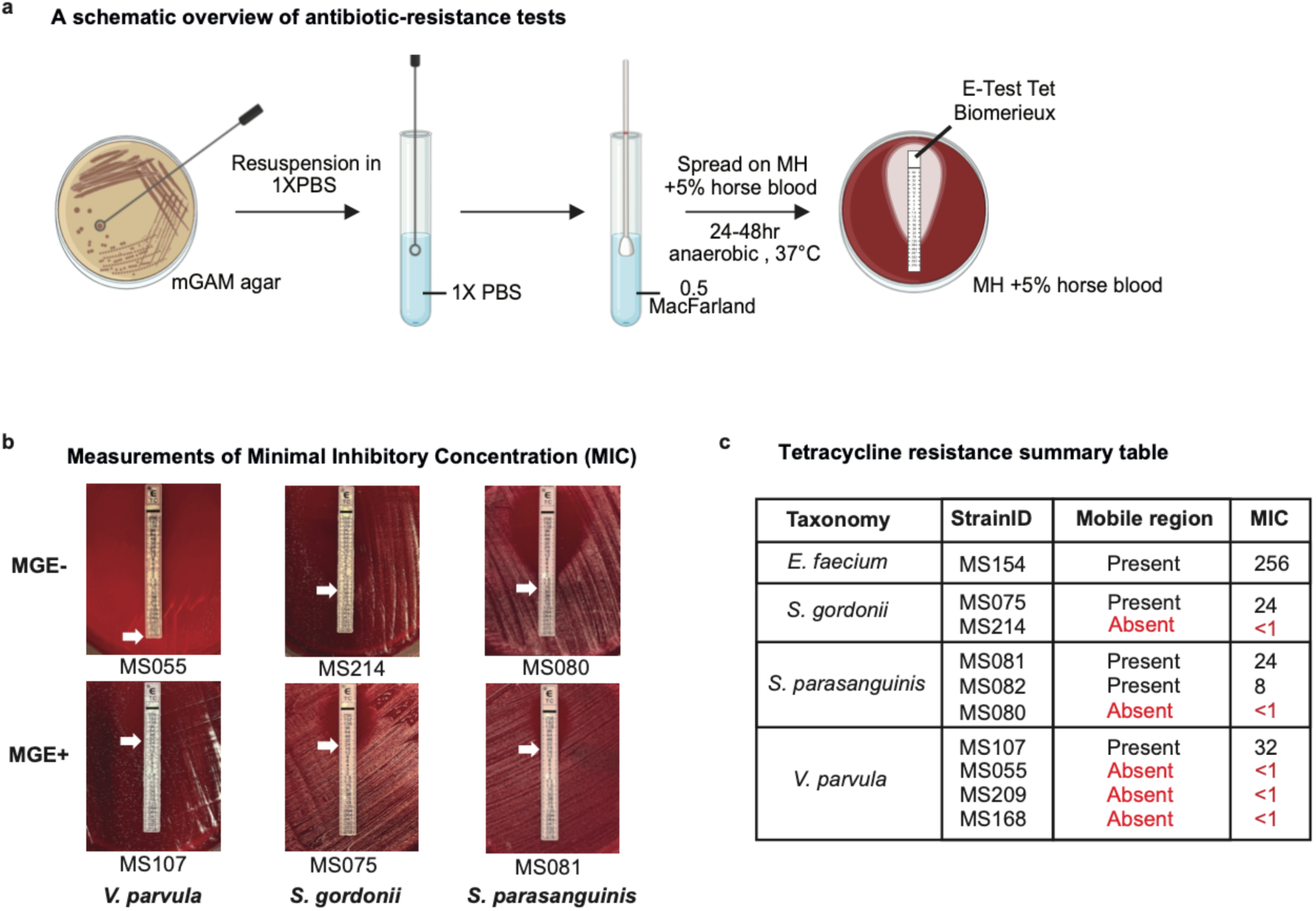
Experimental validation of tetracycline resistance in connection with a mobile element. **(a)** For each strain, several single colonies were picked from an agar plate and resuspended in sterile PBS. The resulting inoculum was then spread onto blood agar plates and an E-test was performed to quantify tetracycline resistance. **(b)** Measurements of Minimal Inhibitory Concentration (MIC) after 24 hours of anaerobic incubation. The MIC for all strains was determined and is indicated by the arrows. Representative examples are shown for species without the mobile element (MGE-) and with the mobile element (MGE+). **(c)** Summary table of tetracycline resistance including the MIC (µg/mL, average of duplicate measurements for each strain).

### MAGraph reveals a nitrate reduction operon duplication event in the *Klebsiella spp.* core genome

A major feature of the MAGraph includes profiling and visualizing bacterial operon structures. We showcase this here with a nitrate reduction (nar) operon, known to facilitate ectopic colonization of oral *V. parvula* in the inflamed gut [14]. First, we searched the MAGraph (IBD_Elinav_2022 cohort) using the functional domains of the *nar* operon genes (**Figure 6a**, left) and retrieved the corresponding subgraphs. The taxonomic assignment of the identified genes was then inferred based on the consensus taxonomy of the nodes within each subgraph (details in method section). For the *V. parvula nar* operon (**Figure 6a**, right) the *narGHJI* genes were localized on the same strand (indicated by black edges) and in close vicinity to each other (intergenic regions of <50bp). Examining their gene neighborhood revealed strain heterogeneity, where a *narLK*-like regulatory system was recovered upstream of the *narGHJI* operon [29] and an S-layer protein was located downstream (ORF3: *SlpA*). The S-layer protein is a class of membrane-associated proteins known to modulate the host immune response [30] [31]. Given that the nitrate reductase complex is membrane-associated, its genomic co-locatisation with S-layer protein genes suggests a potential role in facilitating S-layer assembly, analogous to the involvement of lipopolysaccharides (LPS) in *Caulobacter crescentus* [32]. While the *narGJI* and *narL* genes were highly conserved, where each gene was represented by a single node (i.e. clustered into one gene family). However, *narK* (transporter protein) and *SlpA* (surface layer protein) exhibited greater heterogeneity and were grouped into multiple nodes corresponding to multiple gene families. We used *V. parvula* RNAseq data [33] to analyse gene co-expression. Both *narL* and *narGHJI* were frequently assembled on the same transcript, while *SlpA* was on a different transcript with an upstream neighboring porin gene containing multiple different transmembrane domains (**Figure S3a**). *SlpA* encodes an important cell surface protein in *Clostridium difficile* that mediates bacterial adhesion to human epithelial cells [34]. Overall, this suggests that *V. parvula* likely interacts with the host immune system and attaches to the host cell surface through *SlpA* and adjacent transmembrane proteins.

**Figure 6:**
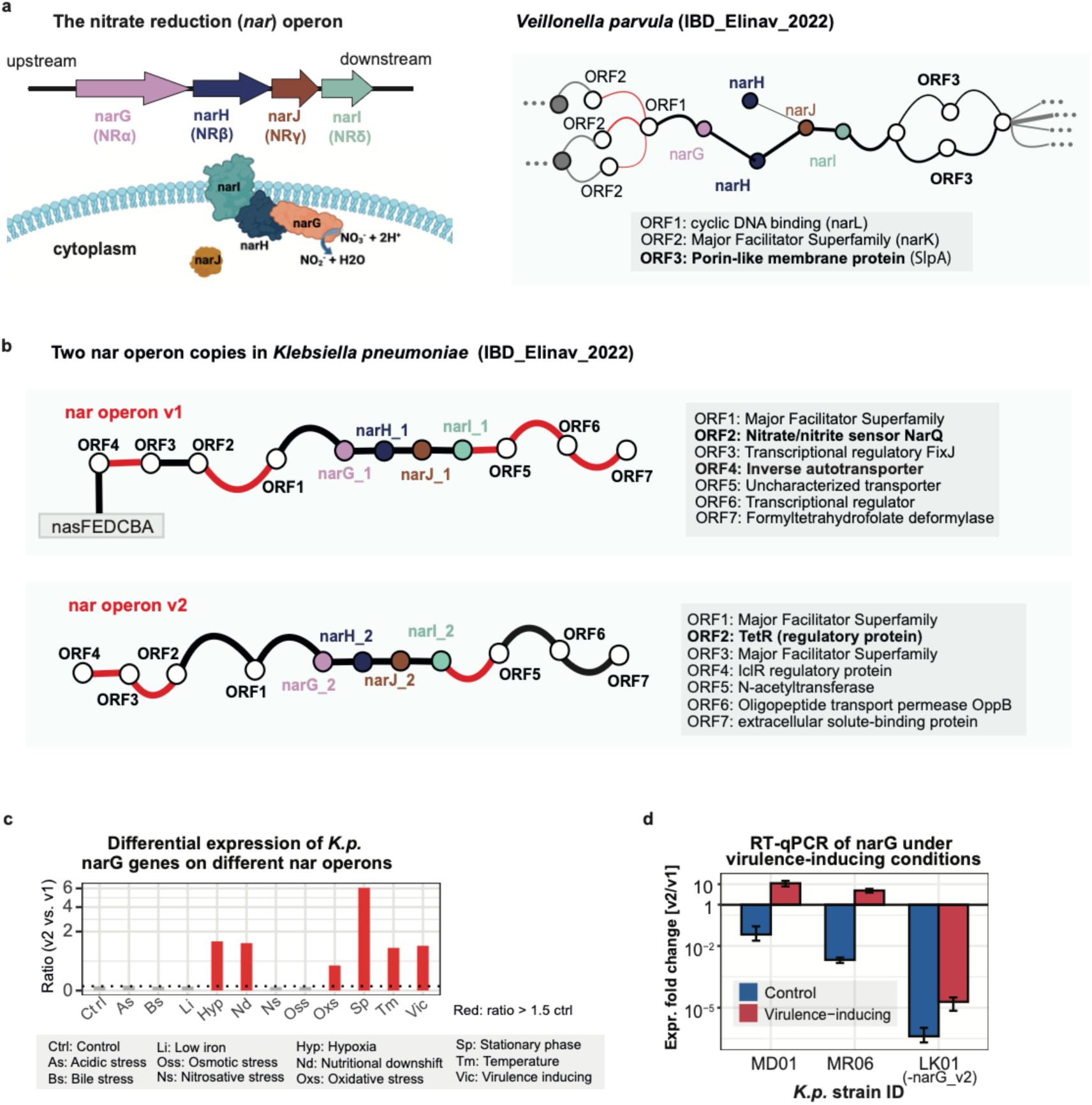
MAGraph reveals differential regulation of duplicated nitrate operons in *K. pneumoniae*. **(a) Left**: Genes in the nitrate reduction *(nar)GHJI* operon including an illustration of the assembly of the nitrate reductase complex: narI is a membrane-associated anchor that binds to *narGH*, while narJ forms a cytosolic chaperon protein, facilitating the formation of the active nitrate reductase complex. **Right**: Subgraph for the *nar* operon in *Veillonella parvula* (IBD_Elinav_2022 cohort). Black edges indicate adjacent genes located on the same strand, while red edges represent genes localized on opposite strands. Straight edges indicate genes in close proximity (mean distance <50 bp), while curved edges indicate greater distances (>50 bp). Edge thickness reflects the frequency of co-localization observed across all contigs. ORF: open reading frame. **(b)** Comparing two *nar* operons in *Klebsiella pneumoniae* (IBD_Elinav_2022 cohort) showed that *nar* operon v1 is located adjacent to a nitrite assimilation operon (*nasFEDCBA*), while *nar* operon v2 is located in close proximity to a Tet Repressor protein (TetR, ∼1.7 kbp upstream). **(c)** Differential regulation of two *K.p. narG* gene copies in response to virulence-inducing conditions. Y-axis shows *narG_v2*:*narG_v1* expression across different culture conditions (x-axis). In the control condition (ctrl), *narG_v1* showed higher expression, while *narG_v2* expression increased in stress and virulence-inducing conditions. **(d)** RT-qPCR validation of the upregulation of the second *narG* gene copy under virulence-inducing condition for clinical *K.p.* isolates. Three strains were tested: MD01 and MR06, which possess both *nar* operons, and LK01, which contains only *nar_v1* as our negative control. The y-axis is the average expression ratio between *nar_v2* and *nar_v1* (n=3, measured using primers targeting *narG_v1* and *narG_v2*). Blue indicates the control group (*K.p.* cultured in LB medium), while red represents virulence-inducing conditions (*K.p.* cultured in LB plus 50% human serum).

Interestingly, for *Klebsiella pneumoniae* two *nar* operon copies were recovered in the MAGraph (**Figure 6b**). *K. pneumoniae* is an IBD-associated pathogen [35] that triggers inflammation in germ-free IL-10 deficient mice [28]. These two *nar* operons were part of the *K. pneumoniae* core genome, evidenced by their linearly correlated abundances indicating co-occurred in all *K.p.* strains rather than representing genes from different strains (**Figure S3b**). Interestingly, these gene copies likely retain similar enzymatic functions as *narGHJI* protein structure predictions were almost identical (qTMscore < 1) despite relatively low sequence similarity (DNA-level similarity ∼75%, **Figure S3c**). While both *nar* operons shared the genes encoding the membrane-associated nitrate reductase (*narGHJI*), their gene neighborhood differed substantially: the first operon (*nar_v1*) was adjacent to a nitrate assimilation operon (*nasFEDCBA*) which converts nitrite to ammonium, while the second operon (*nar_v2*) was located downstream of a transcriptional repressor protein (ORF2, *TetR*) (**Figure 6b**). These distinct gene neighborhoods suggest differential regulation and potentially divergent functions.

### Different *K. pneumoniae nar* operons are linked to differential responses to virulence-inducing conditions

We next investigated the functional implications of this *nar* operon duplication event in *K. pneumoniae* isolates. As gene duplications often lead to functional and regulatory divergence, we tested differences in gene expression under virulence and stress conditions. First, we re-analysed publicly available RNAseq data [36] and showed that *nar_v1* was predominantly expressed under control conditions, while *nar_v2* became the major variant under stress conditions, such as hypoxia, nutritional downshift and virulence-induction (**Figure 6c**). Next, we tested *nar* operon induction by human serum in clinical *K. pneumoniae* isolates. While the first RNAseq dataset was based on a reference strain [36] and provided initial insights, our targeted RT-qPCR approach allowed more precise quantifications of *nar* operon expression and enabled validation across clinically relevant isolates. We analysed *nar* operons in genomes from *Klebsiella* clinical isolates and selected three for further experiments (two with both copies of nar operon, one with only nar_v1). First, serum resistance testing confirmed that all *K. pneumoniae* isolates were able to survive in human serum (**Figure S4d**). Interestingly, *nar_v1* was predominantly expressed under normal growth conditions, whereas *nar_v2* became dominant during virulence induction (**Figure 6d**). Overall, our data indicates that an ancient duplication event led to two *nar* operon copies in *K. pneumoniae*, where the second copy (*nar_v2*) may confer an advantage under virulence-inducing conditions.

We further examined gene evolution dynamics behind this *nar* operon duplication using large-scale comparative genomics. The MAGraph revealed that *Klebsiella oxytoca* also possessed two nar operon copies (**Figure S3d**) with identical gene neighborhoods compared to *K. pneumoniae*, suggesting an ancient duplication event. Sequence alignment supports that the *narG* copies are paralogs as *narG_v1* and *narG_v2* form two clusters across different species (**Figure S3e**). Examining evolutionary duplication and loss events of *narG* across a broader taxonomic range in the *Enterobacteriaceae* family revealed a total of 245 *narG* homologs from 217 different *Enterobacteriaceae* species. Multiple sequence alignment indicates a clear separation into two clades, corresponding to *narG_v1* and *narG_v2*, respectively (sequence id >80% within each clade, **Figure S4a**). An interesting pattern emerged when overlaying the occurrences of *narG* homologs onto a phylogenetic tree constructed from the underlying 217 *Enterobacteriaceae* species genomes (**Figure S4b**): *narG_v1* was more prevalent and found in all species except for certain endosymbionts, while *narG_v2* was less common and only present in species that also contained *narG_v1*. This suggests that *narG_v2* likely derived from a duplication of *narG_v1*. Additional evidence from gene neighborhood analysis further supports this hypothesis: a *tetR* gene was consistently located 1.7 kbp upstream of *narG_v2* across various species. Given that the *tetR* gene was absent from the *nar_v1* neighborhood, this conserved arrangement around *narG_v2* likely reflects a single gene duplication event. We reconstructed the suggested evolutionary events (**Figure S4c**): The common ancestor of *Enterobacteriaceae* species likely contained a single copy of the *nar* operon (*nar_v1*). A duplication event occurred after the divergence of the endosymbiont clade, leading to the emergence of *narG_v2*. The original *narG_v1* was likely lost in the endosymbiont clade shortly after its division, resulting in the absence of the *nar* operon in all endosymbiont *Enterobacteriaceae*. Additionally, *narG_v2* supposedly underwent several gene loss events over time in the non-endosymbiont *Enterobacteriaceae* species. In summary, these findings demonstrate that the MAGraph is a powerful tool for profiling bacterial operons from metagenomic data, providing valuable insights into operon evolution and differential functionality.

## Discussion

Functional microbiome profiling is crucial in order to identify and understand host-microbial mechanisms involved in diseases. In this study, we addressed two major challenges in gene-centric analyses. Firstly, we developed a NoSQL database infrastructure for functional microbiome profiles to facilitate easy and efficient access to the millions of genes profiled with metagenomic sequencing data. Our gene-centric framework MetaGEAR can be applied to any metagenomic dataset of interest, providing a complete workflow from raw reads to the final database. We demonstrated the scalability and capacity of this setup by generating a multi-cohort database comprising more than 300 billion entries profiled from >9000 microbiome samples, which can now be rapidly searched to identify gene signals across multiple cohort studies. Furthermore, in addition to the increased statistical power that this multi-cohort approach offers, it also provides increased taxonomic resolution of individual genes through data integration from multiple cohorts (by propagating gene annotations). The multi-cohort database enabled us to identify IBD and CRC signature genes from specific species that were consistently and robustly associated with disease across studies. Importantly, in the future this data resource can be extended to include additional cohorts and diseases and more datatypes (such as metatranscriptomics and metabolomics data) can be added as additional layers to the database. Secondly, we introduced a graph-based approach (MAGraph) to extract and visualize gene neighborhoods. This offers detailed information about the functional genomic context of individual microbial genes. Importantly, the MAGraph is constructed directly from the gene catalog of the respective metagenomic cohort, which eliminates the need for co-assembly graphs and thus enables graph generations involving hundreds or even thousands of metagenomic samples. As a result, the MAGraph facilitates cohort-level exploration of gene neighborhoods and their respective association with disease.

The MAGraph offers a wealth of information that can be inferred based on examining the genomic context of genes within individual cohorts or across cohorts. This includes profiling of operon structures, strain-level variation of genes, horizontal gene transfer events, and linking mobile elements to their respective bacterial hosts. Furthermore, so far uncharacterized genes and protein domains can be linked to annotated genes and pathways through contextual genomic co-localization. While the majority of the nodes in the MAGraph formed linear structures, a subset of nodes (∼1%) were hyper-connected with >10 connecting edges to other nodes. We found that the majority of these central hubs represented integrase genes that interlinked almost half of all bacterial genes (>49%) into one connected component. Among these hub nodes was a broad-host-range MGE carrying a tetracycline resistance gene that was shared among microbes from diverse phylogenetic lineages. Tetracycline is a widely used broad-spectrum antibiotic with low toxicity; however, its clinical effectiveness has declined due to the increase of antibiotic resistance spreading [37]. We experimentally confirmed that this MGE confers tetracycline resistance in opportunistic pathogens, including *Veillonella*, *Streptococcus*, and *Enterococcus* clinical isolates. Importantly, we observed that other bacterial genera were predicted to contain this MGE, including gut commensals such as *Amedibacterium* and *Erysipelatoclostridium*. Overall, while MGEs are well established as mediators of gene transfer between bacterial species, our findings highlight the critical role of MGEs in shaping the human gut microbiome and underscore the importance of viewing the microbiome as a genetic network, in which horizontal gene transfer facilitates the extensive exchange of genetic resources across species boundaries.

Our study demonstrated the benefits and possibilities of gene-centric analyses based on the MetaGEAR framework; however, there are also limitations that apply. Our gene profiling approach currently relies on the user to provide a particular gene of interest for the search. Next steps in the MetaGEAR development will focus on data-driven approaches for disease-associated gene identification. This will also include the prediction of bacterial operons and biosynthetic gene clusters directly from the MAGraph, thereby extending existing genome- and MAG-based approaches [38] [39] [40] to graph-based representations. This would facilitate robust and accurate predictions including direct visualizations of strain-level variations and disease-associations based on the graph.

In summary, the MetaGEAR framework provides versatile tools and data resources that facilitate gene-centric and functional metagenomic data analyses that will accelerate microbial functional insights into human health and disease.

## Author contribution

S.J. developed and benchmarked the MetaGEAR pipeline, designed and generated the multi-cohort database, identified disease signature genes, showcased the MAGraph framework and wrote the manuscript. W.Y.C. and S.J. developed the MAGraph data object. E.R. designed the pipeline architecture, led the Nextflow implementation, and integrated pipeline modules. E.R., T.E., and S.J. designed and implemented the MAGraph web application. A.C. performed the tetracycline resistance experiments, with advice from S.D., using clinical isolates obtained by D.W. from samples collected by M.St. and M.M. under the supervision of V.C.P. S.D. and S.J. designed *narG* primers. T.R.L. conducted the *nar* operon screening in *Klebsiella* clinical isolates, while L.E. performed the serum resistance assays on these isolates and carried out RT-qPCR experiments, under the supervision of T.S.. A.G. and S.S. conducted protein structure comparisons. T.T.Z. assisted with processing metagenomic datasets. M.Sch. conceptualized the project, acquired funding, supervised all work aspects of the project and wrote the manuscript.

## Conflict of interest

V.C.P. has consulted for Alfasigma S.p.A., Menarini Diagnostics Ltd, AstraZeneca, Emles Bioventures and Resolution Therapeutics, and delivered paid lectures for Norgine Pharmaceuticals Ltd.

## Acknowledgements

This work was supported by grants to MS: Emmy Noether Award (Deutsche Forschungsgemeinschaft; DFG project number 426120468), ERC Starting Grant (European Union, Project: HEROINE - 101039493) and SFB 1371 (DFG project number 395357507.). TS was supported by the Federal ministry of Science of Lower Saxony (project: 11-76251-4658/2022 ZN 4037) and an ERC CoG (as co-PI: project number 865466). Views and opinions expressed are however those of the author(s) only and do not necessarily reflect those of the European Union or the European Research Council. Neither the European Union nor the granting authority can be held responsible for them.

## Method

### Code availability

The MetaGEAR pipeline and MAGraph interface are available through GitHub: https://github.com/schirmer-lab/metagear-pipeline

https://github.com/schirmer-lab/magraph-api

### Details for the metagenomic pipeline for the generation of functional profiles

The pipeline for the generation of functional microbiome profiles is implemented in nextflow and includes four major components. The first part is quality control: raw metagenomic reads are first filtered to remove adapters, low-quality sequences, and human host contamination using the dna_qc command of the MetaGEAR pipeline, which runs TrimGalore (0.6.10) and KneadData (0.10.0). For processing human microbiome datasets in this study, the host reference genome was set to hg37. Updates or alternative host genomes can be configured as well. The second part involves generating a non-redundant gene and protein catalog. Briefly, for each sample, clean reads are *de novo* assembled into contigs using MEGAHIT (v1.2.9). Protein-coding genes are then predicted using Prodigal (v2.6.3) with the “-p meta” flag optimized for metagenomic data. Incomplete genes are filtered out and gene sequences from all samples are merged. The merged gene sequences are clustered using CD-HIT-EST (v4.8.1) to construct a non-redundant gene catalog based on >95% nucleotide sequence identity and >90% coverage. Representative sequences for each gene cluster are subsequently grouped into a protein catalog using CD-HIT (v4.8.1) with amino acid-level >90% identity and >80% coverage. The third part involves abundance profiling and taxonomic annotation of gene families. The abundance of each gene family in each sample is estimated by mapping cleaned reads to the non-redundant gene catalog using Bowtie2 (v2.4.1). Read counts and reads per kilobase per million mapped reads (RPKM) are then calculated from the alignment results using SAMtools (v1.10). Gene families are grouped into metagenomic species pangenomes (MSPs) using MSPminer (v1.1.3), based on co-abundance patterns across samples. The abundance of each MSP is subsequently estimated as the median abundance of its core genes and integrated with MetaPhlAn3 taxonomic profiles based on abundance correlations (as described below). The final part of the pipeline involves protein annotation. Pfam domains are predicted for each representative sequence of the protein families using InterProScan (version 5.47-82.0) and the results are organized into functional domain architectures, capturing both the identified Pfam domains and their sequential order within each protein.

### Integration of reference- and assembly-based profiles

For each MSP and each species predicted by MetaPhlAn3, a linear model was fitted to assess abundance correlation. The mean squared error and R² score were calculated and the best match for a given MSP was determined based on the MetaPhlAn3 species with the highest R² score. An assignment was considered valid for R² > 0.7. If for a given match both the MSP GTDB-Tk annotation and the MetaPhlAn3 species annotation did not match, they were merged and mapped to the next common NCBI taxonomic lineage.

### Evaluation of the taxonomic annotations based on the integrated analysis

The accuracy of taxonomic assignments based on linear associations was evaluated using a subset of MSPs with high-confidence annotations, i.e. those for which the MetaPhlAn3-based assignment agreed with the GTDB-Tk-based assignment at species level. For each of these MSPs, we examined whether the second-best match (i.e. the species with the second-highest R² score) exceeded the threshold of R² > 0.7. Cases where the second-best match surpassed this threshold were considered false positives. Accuracy was then calculated as the percentage of high-confidence MSPs for which the second-best match did not exceed the R² threshold.

To evaluate the detection limits of MSPminer, we used the set of MSPs with high-confidence taxonomic annotations. For each MSP in a given sample, we estimated the number of reads that originated from the corresponding species by multiplying the species’ relative abundance from MetaPhlAn3 with the total number of input reads. These values were binned (bin size = 500). For each bin, we then calculated the empirical detection likelihood as the proportion of samples where MSPminer reported a non-zero abundance. The scatter plot was then generated across all cohorts with the estimated number of species-specific reads on the x-axis and the empirical detection likelihood on the y-axis.

The improvement in taxonomic annotation was evaluated by calculating the mean relative abundance of MSPs that received species-level assignments through the integration with MetaPhlAn3 profiles but were not originally annotated at species-level by GTDB-Tk. This value reflects on average the proportion of the MSP abundance profile that gained species-level resolution through the integration process.

### Data management with the NoSQL database

We provide a script to create a mongoDB database using the output files generated by the pipeline. The mongoDB can be run as a singularity image, which easily facilitates its setup in connection with any server environment. After the database is created, users can query the database using API functions (Python) that we created to extract information, such as gene abundance profile, taxonomic and functional annotations, gene and protein sequences. All scripts are available through github (single_cohort_API.py).

### Establishing a multi-cohort database and identifying disease signature genes

First, we applied our MetaGEAR pipeline to 24 publicly available metagenomic cohorts, comprising 9,053 stool samples from individuals with IBD, CRC, and healthy controls. We then merged all representative sequences from each cohort-level gene catalog to construct a non-redundant multi-cohort gene catalog (MCGC) using CD-HIT-EST (v4.8.1) at >95% nucleotide identity and >90% coverage. These representative gene sequences were translated into protein sequences and clustered into a multi-cohort protein catalog (MCPC) using CD-HIT (v4.8.1) at >90% amino acid identity and >80% coverage. Finally, we organized the MCGC and MCPC cluster information into a MongoDB database and developed a set of Python APIs to support information retrieval. This includes the extraction of abundance profiles, taxonomic classifications, and functional annotations for any set of MCGC or MCPC clusters across all cohorts.

We calculated summary statistics for each MCGC cluster, including the number of IBD, CRC, and healthy cohorts, in which the cluster was identified (denoted as N_IBD_, N_CRC_, N_healthy_). Disease signature genes were identified based on these statistics. In the case of IBD signature genes, we first calculate the frequency fold change in all IBD cohorts FC_IBD_ as:

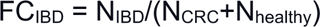

Subsequently, MCGC gene clusters were filtered to obtain gene clusters with N_IBD_>3 and FC_IBD_>10. We selected the top 100 MCGC clusters with the highest FC_IBD_ for further analyses. Importantly, this selection process did not take sample-level disease annotations into account. Instead, candidate gene clusters were required to be consistently detected in IBD cohorts while being absent from healthy populations and other disease cohorts. These signature genes were then examined for disease associations in each cohort. All comparisons were summarised in a volcano plot, where each gene family in each cohort is compared between the disease and control group (Wilcoxon test, mean abundance fold change).

### Construction of a metagenomic assembled graph (MAGraph)

We developed a graph-based data structure, the MAGraph, for exploring and visualising gene neighborhoods based on metagenomic assemblies. In the graph, nodes represent gene families and edges connect pairs of gene families that are found adjacent to each other on the same contig(s). MAGraph operations are supported by a set of Python APIs that interface with the underlying database. A typical workflow begins by selecting a list of gene families, either through sequence homology searches against gene or protein catalogs or by querying for specific functional domains. For each selected gene family, the genomic context is retrieved, including contig IDs, gene coordinates, and strand orientation. Neighboring genes are identified within a user-defined window size on the same contig. These relationships are used to construct an igraph object, where edge properties capture the frequency of co-occurrence, average intergenic distance, and strand information. Functional and taxonomic annotations are then added to the graph as node attributes. Low-confidence edges (i.e. supported by few contigs) are removed based on a user-defined threshold (default = 3). The resulting graph is partitioned into connected components, each representing a subgraph containing the gene(s) of interest. Users can filter these subgraphs based on the number of query-matching nodes. Finally, subgraphs receive taxonomic annotations, which are determined by propagating node-level annotations. Their abundance per sample is estimated based on the median abundance of the nodes.

### MAGraph statistics

The node degree distribution for the MAGraphs was computed by iterating over all gene families and counting the number of directly adjacent nodes (using a window size of 1). This analysis was conducted for all 24 cohorts and the results were aggregated into one barplot. Nodes with a degree greater than 10 were classified as highly connected (hub nodes) and selected for further functional analysis.

To assess graph connectivity, we calculated the size of the largest connected component within each cohort by constructing a graph from all gene families. The connectivity metric was expressed as the fraction of nodes in the largest component divided by the total number of gene families in the cohort.

### Gene Ontology (GO) analysis

The GO enrichment analysis was performed using the topGO package in R (v2.46.0). Briefly, hub gene families in the MAGraphs (degree>10) were selected as the positive set and all gene families assembled from the metagenomic cohorts as the negative set. To assess the statistical significance of GO term enrichment, Fisher’s exact test was applied using the “classic” algorithm. Subsequently, GO terms were ranked according to adjusted p-values.

### Profiling integrases in the MAGraph

First, all integrase-related Pfam domains were used as queries against the multi-cohort protein catalog to identify potential integrases. The matched multi-cohort gene families were then retrieved for downstream analyses. Afterwards, for each gene family, we examined the gene neighborhood (window_size = 10) in each cohort and species associated with the respective integrase were identified based on the taxonomic annotations of the integrase-adjacent genes. We prioritized integrase gene families linked to diverse bacterial hosts for the downstream analysis.

### Profiling a nitrate reduction operon in the MAGraph

The *E. coli nar* operon (*narGHJI*) was used as a query to identify similar nar operon structures in the metagenomic assemblies. For this analysis, we focused on the IBD_Elinav_2022 cohort [28]. The first step involved the prediction of Pfam domains for each query gene from the narGHJI operon using InterProScan (version 5.47-82.0). These domains were then grouped into domain architectures with each group comprising a list of Pfam domains ordered by their appearance in the original query. Next, we searched the cohort protein catalog to identify protein families matching the input functional groups and extracted the corresponding gene families. Subsequently, the corresponding gene neighborhoods for all gene hits were retrieved from the MAGraph (window size = 3, edges were trimmed when supported by <3 contigs). Connected components that contain >2 queries were extracted as subgraphs and annotated by integrating taxonomic and abundance information from the corresponding nodes. Subgraph abundance was estimated by summing up the abundances of all matched genes for each query and taking the median abundance across all queries. Taxonomic annotation of each subgraph was inferred based on the majority consensus of node-level taxonomic assignments.

### Profiling *narG* genes in *Enterobacteriaceae* genomes

We collected 217 reference genomes (selecting the type strain for each species) from the *Enterobacteriaceae* family and built a blastn database. We then used blastn to map the DNA sequences of *narG_v1* and *narG_v2* identified from *K. pneumoniae* against this *Enterobacteriaceae* database. The best match in each genome for each query was selected and subsequently filtered by >80% coverage and >80 sequence identity. Then, all DNA sequences of *narG* homologs were aligned using Clustal Omega [41] with default parameters. The predicted phylogenetic tree was downloaded and visualized using iTOL. Concurrently, the phylogenetic tree of all genomes in the *Enterobacteriaceae* database was constructed using PhyloPhlAn [42] and annotated for present/absence of *narG_v1* and *narG_v2* genes using GraPhlAn (https://huttenhower.sph.harvard.edu/graphlan/).

### Measurement of tetracycline resistance among clinical isolates

Characteristics of the bacterial strains used for tetracycline resistance testing are summarised in **Table 2**. Bacterial growth was performed at 37°C in an anaerobic chamber (Whitley M45, Meintrup DWS Laborgeräte GmbH) at an atmosphere of 5% H2, 5% CO2, and 90% N2. MS055, MS107, MS168 and MS209 were cultivated on Fastidious Anaerobic Agar plates (ThermoFisher, PB0225A). MS081, MS082, MS080, MS075, MS214 and MS254 were cultivated on mGAM agar plates (modified Gifu Anaerobic Medium, Himedia).

Minimal Inhibitory Concentration (MIC) was determined by E-test Tetracycline (TC) (Biomerieux, 423819) on F.A.A (Fastidious Anaerobe Agar) + 5% horse blood. All strains had morphologically similar colonies. A few single colonies were taken from each plate and resuspended in sterile PBS (1X) to reach 0.5 Mac Farland. The inoculum was spread on F.A.A + 5 % horse blood agar plates and dried for 10 minutes. Once the plates were completely dried, an E-test TC was placed on the surface of the agar. Plates were prepared in duplicates for each strain and incubated at 37°C in anaerobic conditions. The values were interpreted using the EUCAST v14.0 clinical breakpoint tables. It should be noted that EUCAST MIC breakpoints are not available for *Veillonella parvula*. Therefore, conclusions were based on inter-species and intra-isolate comparisons as shown in **Figure 5b+c**.

### Differential expression analysis of two *narG* genes in *Klebsiella pneumoniae* isolates

Protein-coding sequences of *Klebsiella pneumoniae* subsp. *pneumoniae* MGH 78578 were downloaded from NCBI. Using the gene sequences of *narG_v1* and *narG_v2* (assembled from the IBD_Elinav_2022 cohort) as queries, homologous genes were identified in the reference genome using blastn (>95% sequence identity and >90% coverage). Their expression profiles (TPM) under different conditions were then obtained from the Pathogenex website (http://www.pathogenex.org/ [36]).

### Serum resistance testing

The survival of *Klebsiella pneumoniae* isolates in 50% human serum was evaluated according to previously described methods [43] with slight modifications. In brief, overnight bacterial cultures were diluted 1:100 in 5 ml of fresh LB medium and incubated at 37 °C shaking at 130 rpm until the OD600 reached 0.1. Bacteria were then pelleted by centrifugation (7,500×g for 5 min at room temperature) and resuspended in 1 mL PBS. Then, an aliquot of 100 µL of the bacterial suspension was added to each well of a 96-well microtiter plate containing 100 µL of human serum (male AB plasma, US origin, Sigma-Aldrich) to give a final serum concentration of 50%. To determine the initial bacterial load, 50 µL of each sample was collected, serially diluted and plated on LB agar plates, followed by overnight incubation at 37 °C. The inoculated microtiter plate was subsequently incubated for 4 hours at 37 °C without agitation. After incubation, bacterial survival was determined by plating serial dilutions on LB agar plates and incubating overnight at 37 °C to determine CFU/ml. Each experiment included a serum-resistant strain (PBIO1289) as a positive control and a serum-susceptible strain (W3110) as a negative control. Serum resistance was quantified as a log2-fold change in CFU/ml after exposure to serum compared to the initial inoculum.

### *Klebsiella pneumoniae* cultivation and virulence-inducing conditions

The growth and stress exposure of *Klebsiella pneumoniae* isolates were performed according to a previously described method with slight modifications [36]. All *Klebsiella* spp. strains were cultured in LB medium at 37°C overnight. Triplicate overnight cultures were diluted 1:100 in 50 ml fresh LB medium and incubated at 37°C with shaking at 180 rpm until an OD600 of 0.5 (exponential phase) was reached. Cultures were then exposed to 50% human serum (male AB plasma, US origin, Sigma-Aldrich) for 1.5 hours with shaking at 180 rpm. Unexposed cultures at exponential growth served as controls for differential expression analysis. To stop bacterial metabolism and stabilize the RNA, 0.5% phenol:ethanol (final concentration) was added to each sample.

### RT-qPCR analysis of the expression of two *nar* operons

One milliliter of triplicate serum-exposed and control cultures was collected with 300 µL of stop solution (95% ethanol, 5% phenol) and snap-frozen in liquid nitrogen. Total bacterial RNA was extracted using the hot phenol method, followed by DNase treatment to remove genomic DNA contamination. RNA concentration was determined by measuring the absorbance at 260 nm using a NanoDrop™ 2000 Spectrophotometer (Thermo Fisher Scientific, USA). For cDNA synthesis, 2 µg of total RNA was reverse transcribed using the RevertAid First Strand cDNA Synthesis Kit (Catalogue #K1622, Thermo Fisher Scientific, USA) according to the manufacturer’s protocol.

The relative expression of two distinct versions of the *nar* operon in *Klebsiella pneumoniae* strains was assessed using RT-qPCR with gene-specific primers (**Table 3**). cDNA synthesized from triplicate serum-exposed and control cultures, as described previously, served as the template for qPCR. Reactions were performed on a CFX96 real-time PCR system (Bio-Rad) using KAPA SYBR® FAST Universal qPCR Master Mix (Kapa Biosystems, #KK4602). Each sample was analyzed in technical triplicates, and PCR-grade water was included as a no-template control to monitor potential contamination. To ensure accurate quantification, cDNA samples used for 16S rRNA quantification were diluted 1:100 prior to amplification.

