## Supplementary figures for "Gene-centric metagenomic analyses reveal microbiome functional insights into diseases"

### Figure S1

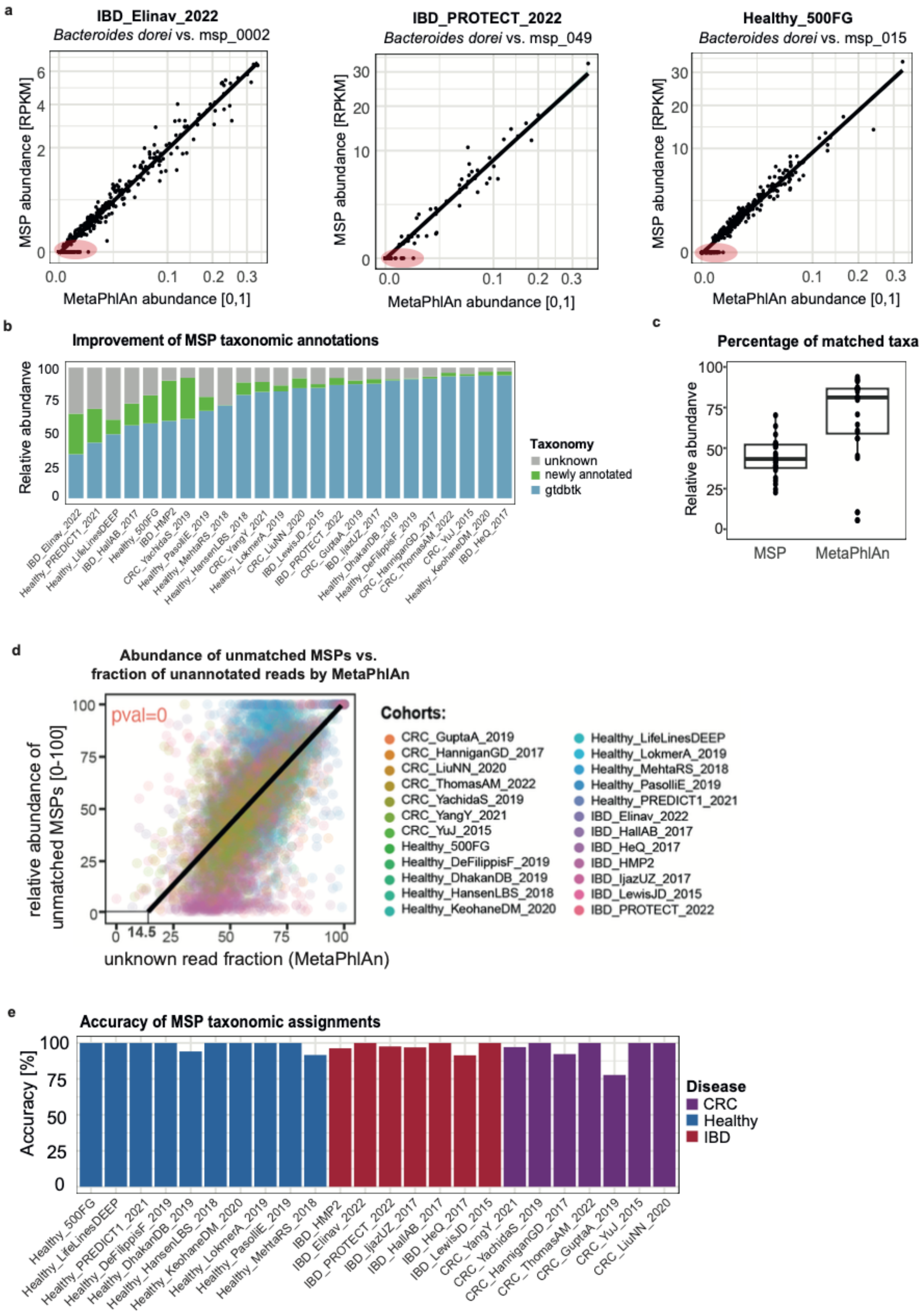
a
IBD_Elinav_2022
Bacteroides dorei vs. msp_0002
IBD_PROTECT_2022
Bacteroides dorei vs. msp_049
Healthy_500FG
Bacteroides dorei vs. msp_015
MSP abundance [RPKM]
MetaPhlAn abundance [0,1]
b
Improvement of MSP taxonomic annotations
Relative abundanve
Taxonomy
unknown
newly annotated
gtdbtk
c
Percentage of matched taxa
Relative abundanve
MSP
MetaPhlAn
d
Abundance of unmatched MSPs vs.
fraction of unannotated reads by MetaPhlAn
pval=0
relative abundance of unmatched MSPs [0-100]
unknown read fraction (MetaPhlAn)
14.5
Cohorts:
CRC_GuptaA_2019
CRC_HanniganGD_2017
CRC_LiuNN_2020
CRC_ThomasAM_2022
CRC_YachidaS_2019
CRC_YangY_2021
CRC_YuJ_2015
Healthy_500FG
Healthy_DeFilippisF_2019
Healthy_DhakanDB_2019
Healthy_HansenLBS_2018
Healthy_KeohaneDM_2020
Healthy_LifeLinesDEEP
Healthy_LokmerA_2019
Healthy_MehtaRS_2018
Healthy_PasolliE_2019
Healthy_PREDICT1_2021
IBD_Elinav_2022
IBD_HallAB_2017
IBD_HeQ_2017
IBD_HMP2
IBD_IjazUZ_2017
IBD_LewisJD_2015
IBD_PROTECT_2022
e
Accuracy of MSP taxonomic assignments
Accuracy [%]
Disease
CRC
Healthy
IBD

**Figure S1: (a)** Linear regression between *Bacteroides dorei* abundances inferred by MetaPhlAn (x-axis) vs. the corresponding metagenomic species pangenomes (MSPs, y-axis). Each subplot represents a different cohort. Highlighted dots (red) are samples where *B. dorei* was only detected in reference-based profiles but not assembly-based profiles. **(b)** The y-axis indicates the cumulative relative abundance of MSPs by category (blue: GTDB-tk annotated; green: improved annotations based on data integration; grey: remaining MSPs with unknown taxonomy) and is averaged across samples for each cohort (x-axis). **(c)** The majority of species detected by MetaPhlAn could be matched to MSPs but not vice versa. The y-axis indicates the cumulative abundance of matched taxa based on our integrative approach (mean value was calculated across all samples for each cohort). Left side: cumulative abundance of MSPs matched to MetaPhlAn species. Right side: cumulative abundance of MetaPhlAn species matched to MSPs. **(d)** The cumulative abundance of MSPs that were not assigned to any MetaPhlAn species explained most of the estimated fraction of unassigned reads in MetaPhlAn profiles. Each point is a sample and the y-axis indicates the cumulative relative abundance of all unmatched MSPs, while the x-axis indicates the estimated fraction of unassigned reads by MetaPhlAn. Colors indicate different cohorts. The fitted linear line indicates a positive correlation (linear regression, $R^2=0.44$, $p<1e^{-308}$). The x-intercept with the correlation line indicates that 14.5% of reads (MetaPhlAn) cannot be assigned to any species even after all MSPs are considered. **(e)** The accuracy (y-axis) of the MSP taxonomic annotation based on the comparison with reference-based taxonomic annotations for each cohort (x-axis). Colors indicate cohort disease focus: Inflammatory bowel disease (IBD), colorectal cancer (CRC) and healthy cohorts.

**Figure S2**

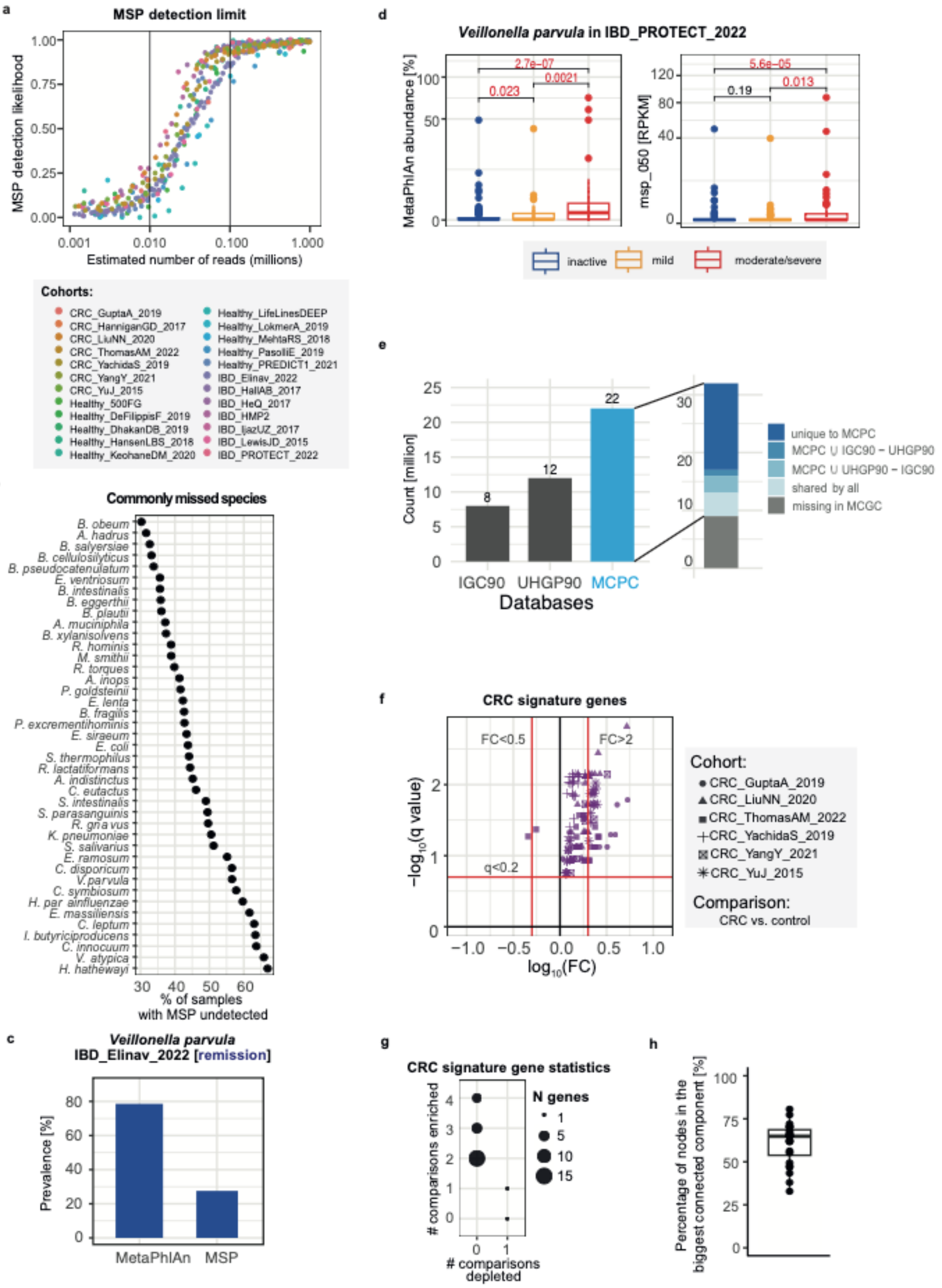

a
MSP detection limit
MSP detection likelihood
Estimated number of reads (millions)
Cohorts:
CRC_GuptaA_2019
CRC_HanniganGD_2017
CRC_LiuNN_2020
CRC_ThomasAM_2022
CRC_YachidaS_2019
CRC_YangY_2021
CRC_YuJ_2015
Healthy_500FG
Healthy_DeFilippisF_2019
Healthy_DhakanDB_2019
Healthy_HansenLBS_2018
Healthy_KeohaneDM_2020
Healthy_LifeLinesDEEP
Healthy_LokmerA_2019
Healthy_MehtaRS_2018
Healthy_PasolliE_2019
Healthy_PREDICT1_2021
IBD_Elinav_2022
IBD_HallAB_2017
IBD_HeQ_2017
IBD_HMP2
IBD_IjazUZ_2017
IBD_LewisJD_2015
IBD_PROTECT_2022
b
Commonly missed species
% of samples with MSP undetected
c
Veillonella parvula
IBD_Elinav_2022 [remission]
Prevalence [%]
MetaPhlAn
MSP
d
Veillonella parvula in IBD_PROTECT_2022
MetaPhlAn abundance [%]
msp_050 [RPKM]
inactive
mild
moderate/severe
e
Count [million]
IGC90
UHGP90
MCPC
Databases
unique to MCPC
missing in MCGC
shared by all
f
CRC signature genes
FC<0.5
FC>2
q<0.2
−log10(q value)
log10(FC)
Cohort:
CRC_GuptaA_2019
CRC_LiuNN_2020
CRC_ThomasAM_2022
CRC_YachidaS_2019
CRC_YangY_2021
CRC_YuJ_2015
Comparison:
CRC vs. control
g
CRC signature gene statistics
### comparisons enriched
### comparisons depleted
N genes
h
Percentage of nodes in the biggest connected component [%]

**Figure S2: (a)** Empirical estimation of the MSP detection limit. Each dot represents a MSP and the x-axis indicates the estimated number of reads per MSP. The y-axis represents the likelihood of detecting this MSP (details in method section). The vertical line at x=0.01 represents the lower empirical threshold, below which an MSP is rarely detected. The line at x=0.1 marks the upper empirical threshold, above which an MSP is consistently detected. **(b)** Summary of species with underestimated prevalence in assembly-based approaches. The x-axis indicates the frequency across all samples and cohorts for each species, which was detected by MetaPhlAn but not present in the MSP profiles. **(c)** Comparing detection sensitivity of *Veillonella parvula* between reference-based (MetaPhlAn, left) and assembly-based (MSP, right) approaches among remission patients in an IBD cohort (IBD_Elinav_2022). MetaPhlAn detected *Veillonella parvula* in 78.6% of remission patients, whereas assembly-based *V.p.* MSPs were only detected in 27.6% of samples from patients in remission, emphasizing the challenge of detecting and assembling IBD-associated species during disease remission. **(d)** Analogous comparison of detection sensitivity for different disease severity groups in an IBD cohort (IBD_PROTECT_2022). MetaPhlAn detected more lowly abundant *Veillonella parvula* signals in mild disease compared to the assembly-based approach, showing a significant increase of *V.p.* in mild compared to inactive disease (p: 0.023, Wilcoxon). **(e)** Comparison of our multi-cohort protein catalog (MCPC) with other publicly available gene catalogs, including the Unified Human Gastrointestinal Protein (UHGP) and Integrated non-redundant gene catalog (IGC). Genes from each catalog were regrouped at the same sequence similarity threshold as our MCPC (>90% identity, >80% coverage, indicated as UHGP90 and IGC90). The y-axis indicates the total number of protein families. **(f)** Volcano plot of colorectal cancer (CRC) signature genes, indicating significance on the y-axis (-log10 transformed q-values, Wilcoxon, Benjamini-Hochberg correction) and fold change (FC) on the x-axis (log10 transformed, mean FC: CRC vs healthy). Red lines indicate thresholds for significance and shapes indicate cohort. **(g)** Statistics of CRC signature genes: indicating the number of comparisons showing enrichment of signature genes (y-axis) versus depletion (x-axis). Point size indicates the number of genes belonging to each category. Almost all CRC signature genes are consistently enriched in multiple CRC cohorts. **(h)** Fraction of nodes contained in the largest connected component of the MAGraph for each cohort.

### Figure S3

**a**

**Gene neighbourhood of *V. parvula* nar operon**

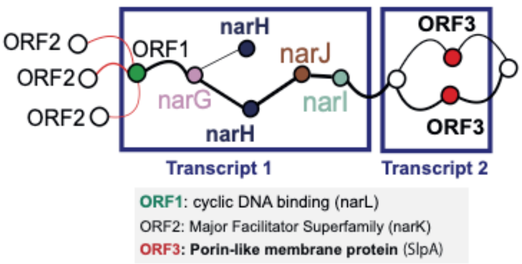


**b**

**Two nar operon copies in *K. pneumonia***

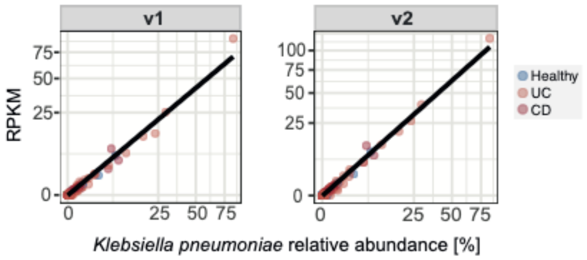


**c**

**Protein structure comparisons between two nar *K. pneumonia* operon copies**

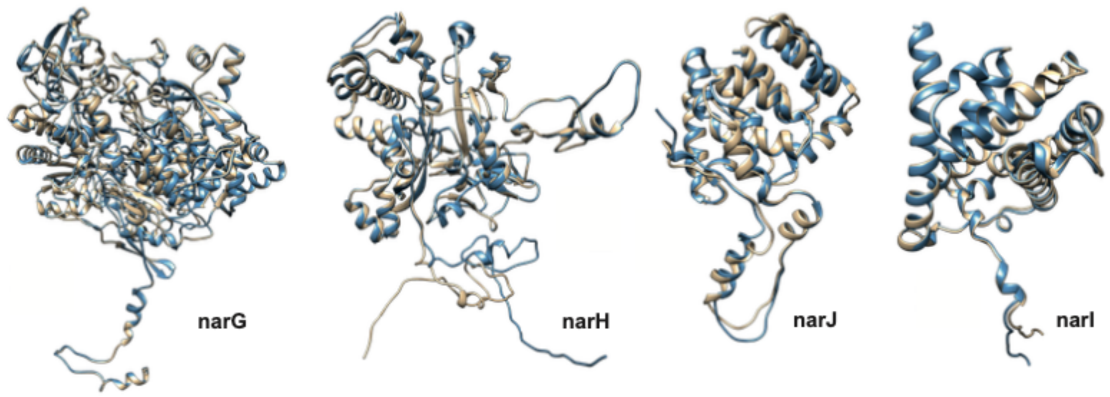


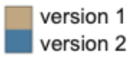


| | DNA | Protein sequence | | Protein structure | |
|---|---|---|---|---|---|
| | identity | identity | similarity | rmsd | qtmscore |
| **narG** | 77% | 79% | 89% | 1.98 | 0.91 |
| **narH** | 79% | 80% | 89% | 6.27 | 0.92 |
| **narJ** | 61% | 54% | 68% | 0.45 | 1.00 |
| **narI** | 75% | 75% | 89% | 1.04 | 0.98 |

**d**

**Two nar operon copies in *Klebsiella oxytoca* (IBD_Elinav_2022)**

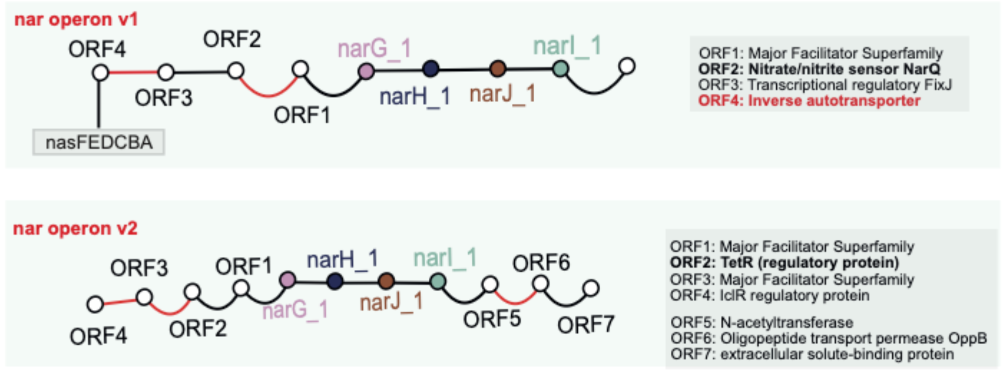


**e**

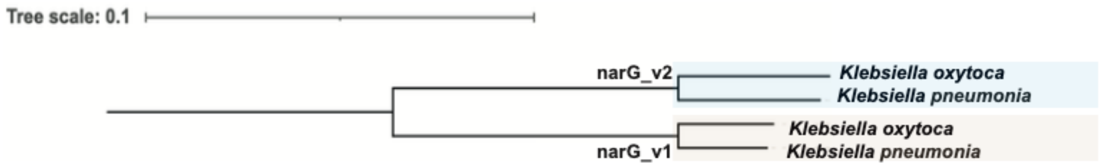

**Figure S3:** **(a)** Integrating *Veillonella parvula nar* operon gene expression data [1] with metagenomic information on gene neighborhood. Blue boxes indicate genes recovered from a single transcript (assembled RNAseq data). The *narGHJI* genes and an anaerobic regulatory protein (ORF1) were recovered from the same transcripts. **(b)** *Klebsiella pneumoniae* has two copies of the *nar* operons in its core genome. The y-axis indicates the abundance of *nar* operons v1 (left) and v2 (right), while the x-axis indicates *K.p.* relative abundance (MetaPhlAn). Both *nar* operons show a linear correlation with *K. pneumoniae* abundance, suggesting that both are part of the core genome. **(c)** The two *nar* operons in *Klebsiella pneumoniae* exhibit highly similar protein structures. The *narGHJI* genes from both operons were compared at gene sequence, protein sequence, and protein structure levels (see table for details). For sequence comparisons, identity refers to the percentage of identical residues in the aligned regions, while similarity accounts for amino acids with comparable physiological properties. For protein structure comparisons, we calculated the q-TMscore (TM-score normalized by query length) and the root mean square deviation (RMSD). Genes compared here are from *K. pneumonia nar* operons profiled in the IBD_Elinav_2022 cohort. **(d)** Two copies of the *nar* operon were also identified in *Klebsiella oxytoca* and their genomic neighborhood was extracted from the MAGraph (IBD_Elinav_2022). **(e)** Phylogenetic analysis of *narG* genes from *K. pneumoniae* and *K. oxytoca*. The *narG_v1* and *narG_v2* belong to *nar* operon v1 and v2, respectively. Multiple sequence alignment based on protein sequences suggests that *narG_v1* and *narG_v2* are paralogs resulting from an ancient duplication event.

#### Figure S4

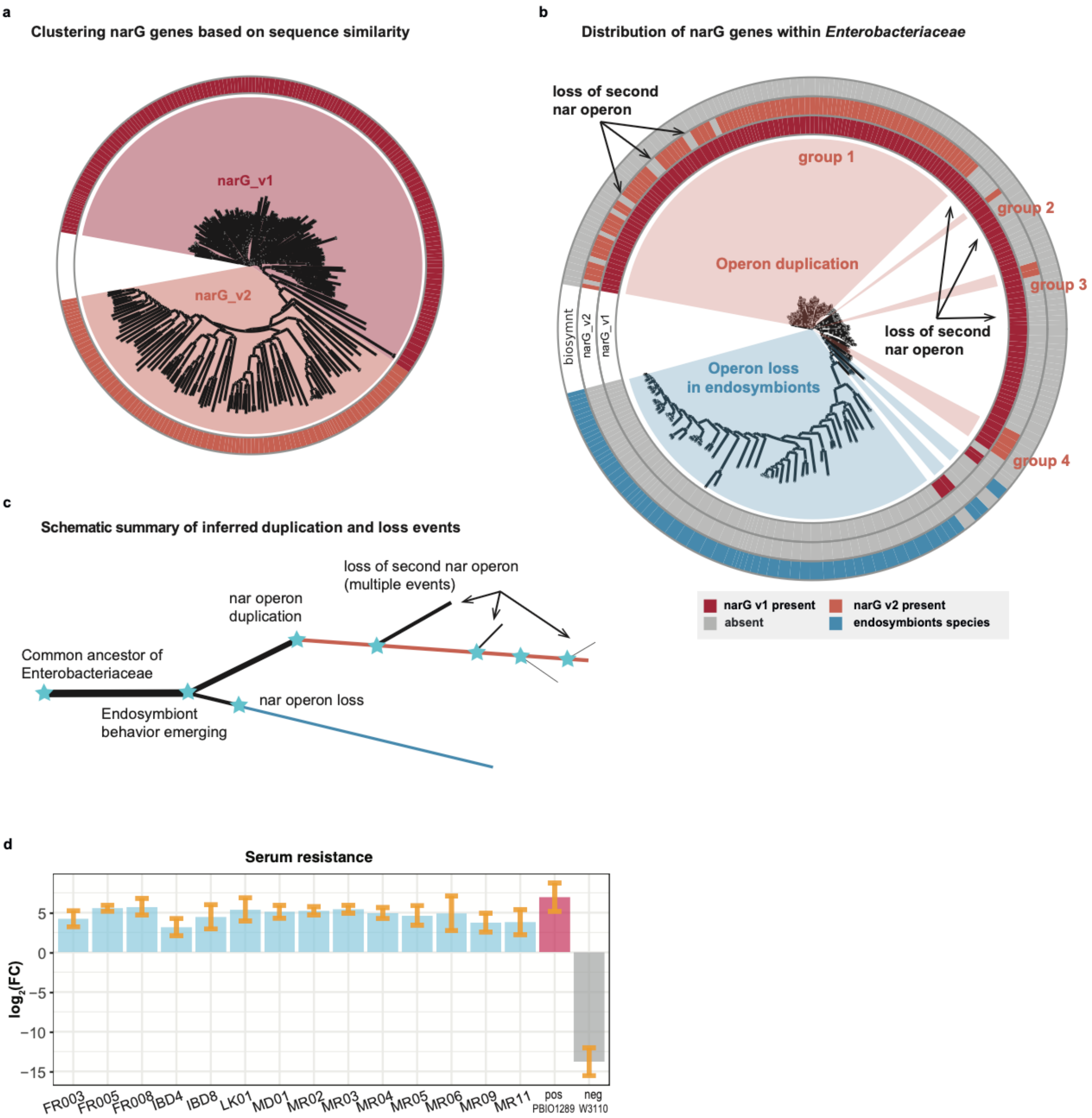

a
Clustering narG genes based on sequence similarity
narG_v1
narG_v2
b
Distribution of narG genes within *Enterobacteriaceae*
loss of second nar operon
group 1
group 2
group 3
group 4
Operon duplication
loss of second nar operon
Operon loss in endosymbionts
biosymnt
narG_v2
narG_v1
narG v1 present
narG v2 present
absent
endosymbionts species
c
Schematic summary of inferred duplication and loss events
loss of second nar operon (multiple events)
nar operon duplication
Common ancestor of Enterobacteriaceae
Endosymbiont behavior emerging
nar operon loss
d
Serum resistance
log2(FC)
5
0
−5
−10
−15
FR003
FR005
FR008
IBD4
IBD8
LK01
MD01
MR02
MR03
MR04
MR05
MR06
MR09
MR11
pos PBIO1289
neg W3110

**Figure S4: (a)** The phylogenetic tree was built based on 245 *narG* homologs (>80% amino acid similarity and >80% coverage) from 217 *Enterobacteriaceae* genomes. The color highlights two major clades, representing *narG_v1* and *narG_v2*. **(b)** Phylogenetic tree of these *Enterobacteriaceae* containing *narG* genes (PhyloPhlAn, using type strain genomes). The two inner rings indicate the presence and absence of *narG_v1* and *narG_v2*. The outer ring highlights species with endosymbiont behavior (or candidatus), which form a separate cluster. Black arrows highlight cases of loss events of *narG_v2*. **(c)** A schematic summary of the hypothesised duplication and deletion events for the *nar* operon. The common ancestor of the *Enterobacteriaceae* family likely had a single nar operon copy. Shortly after the division of the endosymbiont clade, a duplication event in the major clade resulted in the second *nar* operon copy located in the vicinity of the Tet Repressor (*TetR*) gene. This was likely followed by multiple gene loss events leading to speciation of the main clades. **(d)** Serum resistance assay for *Klebsiella pneumoniae* clinical isolates. Strain names are indicated on the x-axis, where *E. coli* strain PBIO1289 is used as positive control and *E. coli* strain W3110 is the negative control.
